# Protein-guided RNA barcoding links transcriptomes to synaptic architecture

**DOI:** 10.64898/2026.02.26.705527

**Authors:** Amanda Urke, Michael-John Dolan, Jonah Silverman, Michael Kim, Juan Pineda, Samantha Garcia, Judy Luu, Alex Buckley, Vipin Kumar, Binhui Zhao, Ken Chan, Naeem Nadaf, Karol S. Balderrama, Don B. Arnold, Beth Stevens, Benjamin E. Deverman, Evan Z. Macosko

**Author notes:** Equal contributions (Amanda Urke, Michael-John Dolan, Jonah Silverman, Michael Kim).

## Abstract

Mammalian brain function relies on the precise synaptic architecture of diverse cell types, yet scalable methods for linking a neuron’s transcriptomic profile to its neuroanatomy remain limited. We present Synapse-seq, an *in vivo* strategy in which cell-identifying barcoded mRNAs are routed to subcellular compartments via targeting proteins and detected by single-cell and spatial genomics. Using AAV delivery for minimal perturbation of gene expression, we directed barcodes to presynaptic terminals (via synaptophysin) in four distinct circuits, or to postsynaptic sites (via nanobodies to endogenous PSD95) of hippocampal excitatory neurons. In the mouse primary visual cortex, presynaptic Synapse-seq recovered known long-range projections and discovered cortical layer subtypes with distinct thalamic innervation. In the anterior cortex, we elucidated simple topographic rules of corticostriatal innervation: intratelencephalic neurons followed a continuous depth-to-target gradient, while extratelencephalic neurons exhibited striatal collaterals that spatially correlated with medullary innervation. Finally, postsynaptic barcoding of excitatory neurons revealed cell type-specific variation in dendritic architectures across and within hippocampal subfields. These data establish Synapse-seq as a versatile, genomics-based approach for the integrated definition of molecular identity and synaptic organization across mammalian brains.

---

Advances in high-throughput single-cell profiling have revolutionized our ability to define cell types across the mammalian nervous system. Recent applications of these technologies have revealed extraordinary molecular variation among neurons^1–5^, yet a critical challenge remains: how to functionally interpret these molecular definitions^6^. Beyond their molecular signatures, neurons are also distinguished by the spatial distribution of their presynaptic and postsynaptic arbors^7–10^, which can vary significantly across, and within, transcriptionally defined subtypes^11,12^. Linked measurements of neuronal anatomy and molecular identity will help elucidate how neurons specialize to carry out diverse brain functions.

Several recent methods have begun to bridge this gap. Retrograde adeno-associated virus (AAV)-based strategies like Retro-seq^13,14^, Epi-retro-seq^15^, VECTORseq^16^ and MERGEseq^17^ robustly map long-range projections prior to single-cell profiling, yet the diffusion of injection material limits their resolution and precludes the characterization of fine-grained spatial topographies. Barcoded transsynaptic tracing with rabies virus can identify pairs of connected cells^18,19^, but neuroinflammatory responses^20^, the need for pseudotyping to control viral spread, and a restricted number of starter cells^21,22^ hinder scalability and routine application. MAPseq uses Sindbis virus to fill axons with RNA barcodes^23^, which, when combined with targeted *in situ* sequencing of selected neuronal genes^24^, can provide a powerful framework for multiplexed presynaptic projection mapping. Yet, Sindbis infection can alter host gene expression and cell health^25^, limiting survival time of injected animals and the ability to overlay these maps onto unperturbed, whole transcriptome profiles defined by standard single-cell atlases.

To address these challenges, we developed Synapse-seq, an AAV-based, protein-guided RNA trafficking platform for highly multiplexed mapping of cell type-specific neural circuits *in vivo*. Barcoded, polyadenylated viral transcripts are routed to a subcellular compartment specified by the endogenous localization pattern of the selected protein targeting domain, for example synaptophysin for labeling presynaptic projections or a nanobody to endogenous PSD95 to direct barcodes towards excitatory postsynapses. These trafficked copies of the cell-identifying barcodes are detected in target sites of interest with bulk and spatial transcriptomics assays and then matched back to the corresponding unbiased whole transcriptome profiles assayed by single-nucleus or spatial transcriptomics at the injection site. Delivering both Synapse-seq components via AAV confers two key advantages: the relatively low immunogenicity of AAV^26^ offers stable, long-term expression with minimal perturbation of endogenous gene-expression profiles, while its popularity as a tool for genetic manipulation of neural circuits supports future innovations. Here, we applied presynaptic Synapse-seq to characterize cell type-specific projections from the primary visual cortex and anterior cortex. Separately, we used the postsynaptic trafficking system to resolve heterogeneity in both gene expression and dendritic morphology across multiple spatial axes of the hippocampus.

## Labeling synaptic compartments with a targeting protein and mRNA barcode

We developed a two-component, AAV-compatible system for the modular targeting of barcoded mRNA transcripts to protein-specified subcellular locations in neurons. Component 1 encodes mScarlet with untranslated regions containing PP7 RNA stem loops, a stability-enhancing pseudoknot^27^, and a 32-nucleotide barcode immediately upstream of a polyadenylation signal (Fig. 1a, Extended Data Fig. 1a). Component 2 encodes a tandem-dimer of the PP7 coat protein (tdPCP) to specifically bind Component 1’s RNA stem loops^28^, fused to a targeting domain that localizes to specific neuroanatomical sites (e.g. synaptophysin for presynaptic terminals^29^, or nanobodies against postsynaptic proteins^30^), and a GFP tag for visualization. We reasoned that binding between tdPCP and PP7 sites would traffic the barcoded transcript according to the localization pattern specified by the targeting protein.

**Fig. 1:**
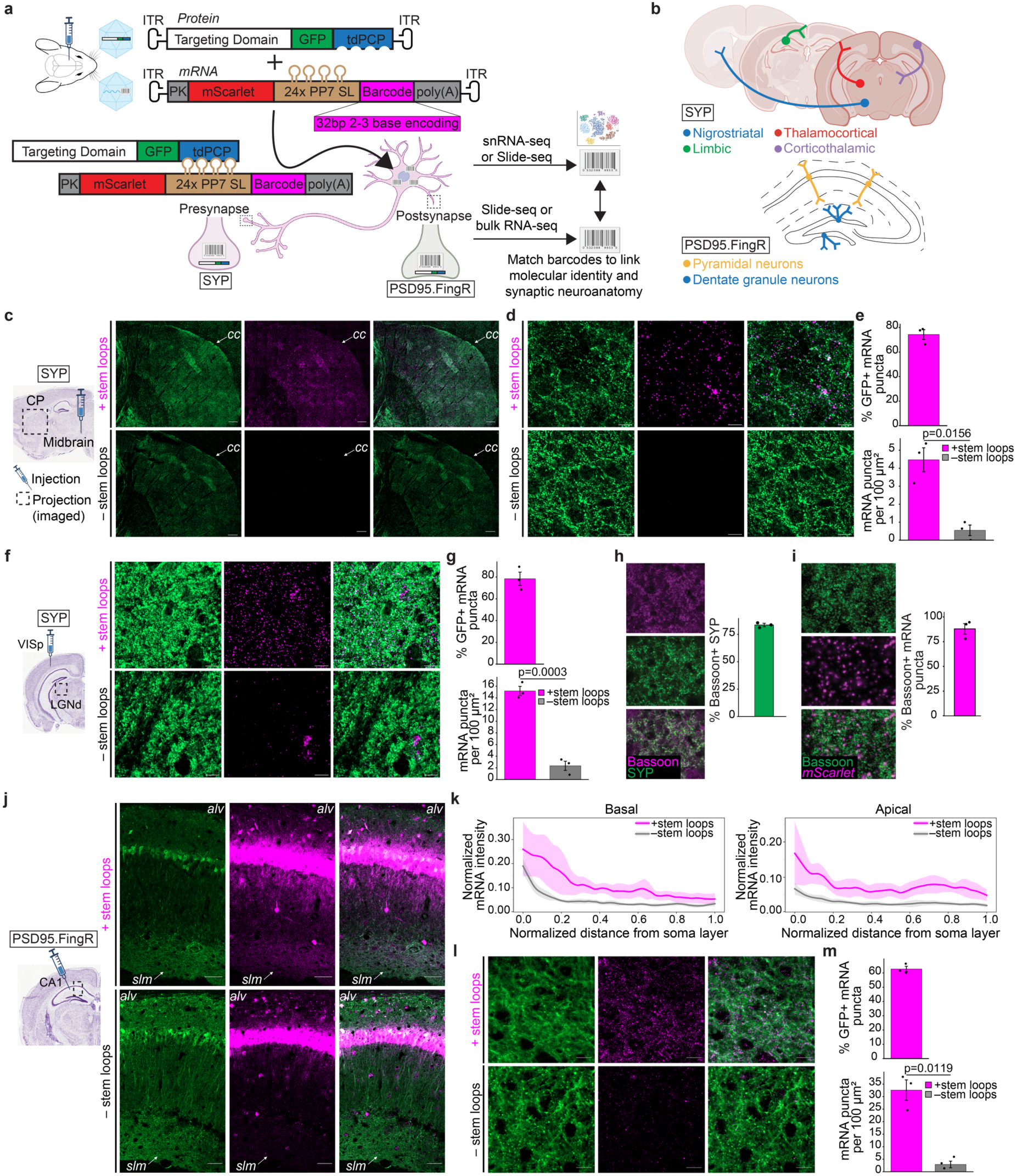
*In vivo* delivery of cell-identifying mRNA barcodes to pre- or postsynaptic compartments. **a,** Workflow for subcellular targeting of barcoded mRNAs. Abbreviations: ITR, inverted terminal repeats; PK, pseudoknot; SL, stem loops; poly(A), hGH polyadenylation signal. **b**, Schematic of cell types and targeting systems (presynaptic: synaptophysin, SYP; postsynaptic: PSD95.FingR). **c**,**d**, SYP-guided labeling of nigrostriatal projections. SYP protein (green, anti-GFP IHC) and *mScarlet* mRNA (magenta, RNAscope smFISH) imaged in the CP at 20x (**c**; scale bars 200 μm; cc, corpus callosum) and 63x (**d**; scale bars 10 μm) with (top) or without (bottom) PP7 stem loops. **e**, Specificity (% mRNA colocalized with GFP, top) and density (puncta per 100 μm^2^, bottom) of mRNA labeling in CP (*n*=3 mice per condition; mean ± s.e.m.; t=5.39, p=0.0156, df=2.75). **f**, SYP-guided labeling of corticothalamic projections. SYP protein and mRNA imaged in the LGd at 63x (scale bars 10 μm). **g,** Specificity and density quantification as in **e** of mRNA labeling in LGd (*n*=3 mice per condition; mean ± s.e.m.; *t*=11.57, *p*=0.0003, df=3.99). **h,i**, Colocalization of anti-Bassoon IHC with SYP targeting protein (**h**, green, native GFP) or *mScarlet* mRNA (**i**, magenta, RNAscope smFISH) imaged at 63x (left) and quantified (right, *n*=3 mice; mean ± s.e.m.). **j-l**, PSD95.FingR-guided labeling of CA1 basal/apical layers. Targeting protein (green) and mRNA (magenta, HCR smFISH) imaged as 63x tile (**j**; scale bars 50 μm; alv, alveus; SLM, stratum lacunosum-moleculare) or 63x ROI of distal apical layer (**l**; scale bars 10 μm) with (top) or without (bottom) stem loops. **k,** mRNA fluorescence intensity vs. distance from soma, normalized to soma layer intensity (*n*=3 mice per condition, 3 measurements/mouse; shaded 95% confidence interval). **m,** Specificity and density quantified as in **e** of mRNA labeling in distal apical layers (*n*=3 mice per condition; mean ± s.e.m.; *t*=6.92, *p*=0.0119, df=2.42).

We first tested this system in primary cortical neurons *in vitro*. When both components were co-transfected, single molecule fluorescent *in situ* hybridization (smFISH) for *mScarlet* showed robust redistribution of the mRNA transcript to the intended synaptic sites: the synaptophysin (SYP) targeting protein drove presynaptic localization, while a nanobody to gephyrin (GPHN.FingR) or PSD95 (PSD95.FingR^30^) resulted in postsynaptic localization (Extended Data Fig. 1b-d). Using PSD95.FingR, we verified that dendritic trafficking was dependent on the presence of stem loops in the mRNA transcript, and concluded that the tdPCP-PP7 system trafficked more efficiently than an alternative tdMCP-MS2 system (Extended Data Fig. 1e,f).

A primary potential application of our technology is the routine, non-toxic mapping of long-range presynaptic projections *in vivo*. We therefore asked whether our trafficking system could be applied to projection neurons possessing different axonal architectures, neurotransmitter identities, and neurodevelopmental origins. Using SYP as our targeting protein, we tested four long-range circuits: nigrostriatal dopaminergic projections from the midbrain to the caudoputamen (CP), thalamocortical projections from the ventral posteromedial nucleus (VPM) to the somatosensory cortex (SSp), hippocampal projections from the CA1 to the retrosplenial cortex (RSC), and corticothalamic projections from the primary visual cortex (VISp) to the lateral geniculate nucleus (LGd) (Fig. 1b).

Following stereotactic delivery of both AAV-packaged Synapse-seq constructs to each injection site of interest, we observed extensive labeling of presynaptic projection territories with both the GFP-tagged SYP targeting protein and mRNA transcripts, detected by smFISH (Fig. 1c-i, Extended Data Fig. 1g-o and Methods). The trafficking of mRNA transcripts was specific to the targeting protein distribution (74-82% of mRNA puncta colocalized with GFP-tagged SYP) and dependent on the presence of PP7 stem loops (6-16-fold enrichment in density of mRNA puncta relative to stem-loop-negative controls) (Fig. 1e,g, Extended Data Fig. 1i,l). In the corticothalamic circuit, we assessed the stability of presynaptic trafficking from 1 to 4 weeks of viral incubation and observed no change in the specificity or density of mRNA labeling over time (Extended Data Fig. 1n-o). Finally, to characterize presynaptic labeling sensitivity, we co-stained the LGd of VISp-transduced mice for the presynaptic scaffold protein Bassoon and found that most SYP targeting protein (83.5%) and mRNA transcripts (88%) colocalized with the marker (Fig. 1h,i).

By exchanging SYP for PSD95.FingR, we explored the capability of our system to map local distributions of postsynapses rather than long-range presynaptic projections. We injected the postsynaptic targeting system into the hippocampus, where the spatial separation of the pyramidal neuron somas from their dendritic branches is ideal for visualizing and quantifying mRNA trafficking. mRNA transcripts, detected by smFISH, accompanied the postsynaptic targeting protein across the full length of basal and apical synaptic layers of the CA1, as evidenced by sustained mRNA fluorescence intensity that was significantly attenuated in the absence of PP7 stem loops (Fig. 1j-k). In the distal apical layer, mRNA labeling was specific (63% of mRNA puncta colocalized with GFP-tagged PSD95.FingR) and stem-loop-dependent (11-fold enrichment in density of mRNA puncta relative to stem-loop-negative controls) (Fig. 1l-m). Collectively, these experiments demonstrate the modularity of our approach for selectively delivering mRNA transcripts to locations within a neuron, as specified by the endogenous targeting of selected proteins.

## Validation of presynaptic Synapse-seq in a canonical corticothalamic circuit

We next combined our presynaptic targeting system with single-nucleus RNA-seq (snRNA-seq) and either bulk or spatial transcriptomics (Slide-seq^31,32^) to match presynaptic sites from long-range projection neurons to their source cells. We focused first on VISp, a region with well-characterized long-range connectivity^14^, to validate the accuracy of our barcoding approach.

After injecting presynaptic Synapse-seq into VISp, we performed snRNA-seq at the injection site alongside targeted amplification and sequencing of the barcoded viral mRNA transcripts (VTs) from the same nuclei (Methods). First, to measure whether Synapse-seq AAV transduction alters native gene expression, we compared our nuclei to a whole-brain single-cell RNA-seq reference atlas^2^ (Extended Data Fig. 2a-c and Methods). Relative to non-transduced cells, and in contrast to Sindbis-transduced cells^24^, Synapse-seq-transduced cells showed no loss of cell-type mapping confidence and retained high marker gene expression concordance (Extended Data Fig. 2d-h), indicating minimal transcriptional perturbation.

The label transfer of our VISp-transduced snRNA-seq dataset produced hierarchical assignments across both neuronal and glial populations (Extended Data Fig. 3a and Methods). Among long-range projecting excitatory neurons, we recovered all major cortical subclasses: intratelencephalic-projecting (IT) cells (L2/3, L4/5 and L6); layer 5 pyramidal tract extratelencephalic neurons (L5 ET); layer 5 near-projecting neurons (L5 NP); and layer 6 corticothalamic neurons (L6 CT) (Extended Data Fig. 3b,c). Across all transduced cell types, we detected a mean of 6.9 unique VTs per nucleus, with modest subclass variation (Extended Data Fig. 3d-f). To map the presynaptic projections of these neurons, we sampled VTs from canonical VISp targets by performing Slide-seq (thalamus, superior colliculus) or bulk RNA-seq (contralateral VISp, striatum, and pons) (Fig. 2a and Methods). In total, after QC (Methods), we recovered 16,968 nuclei with joint RNA and VT expression data.

**Fig. 2:**
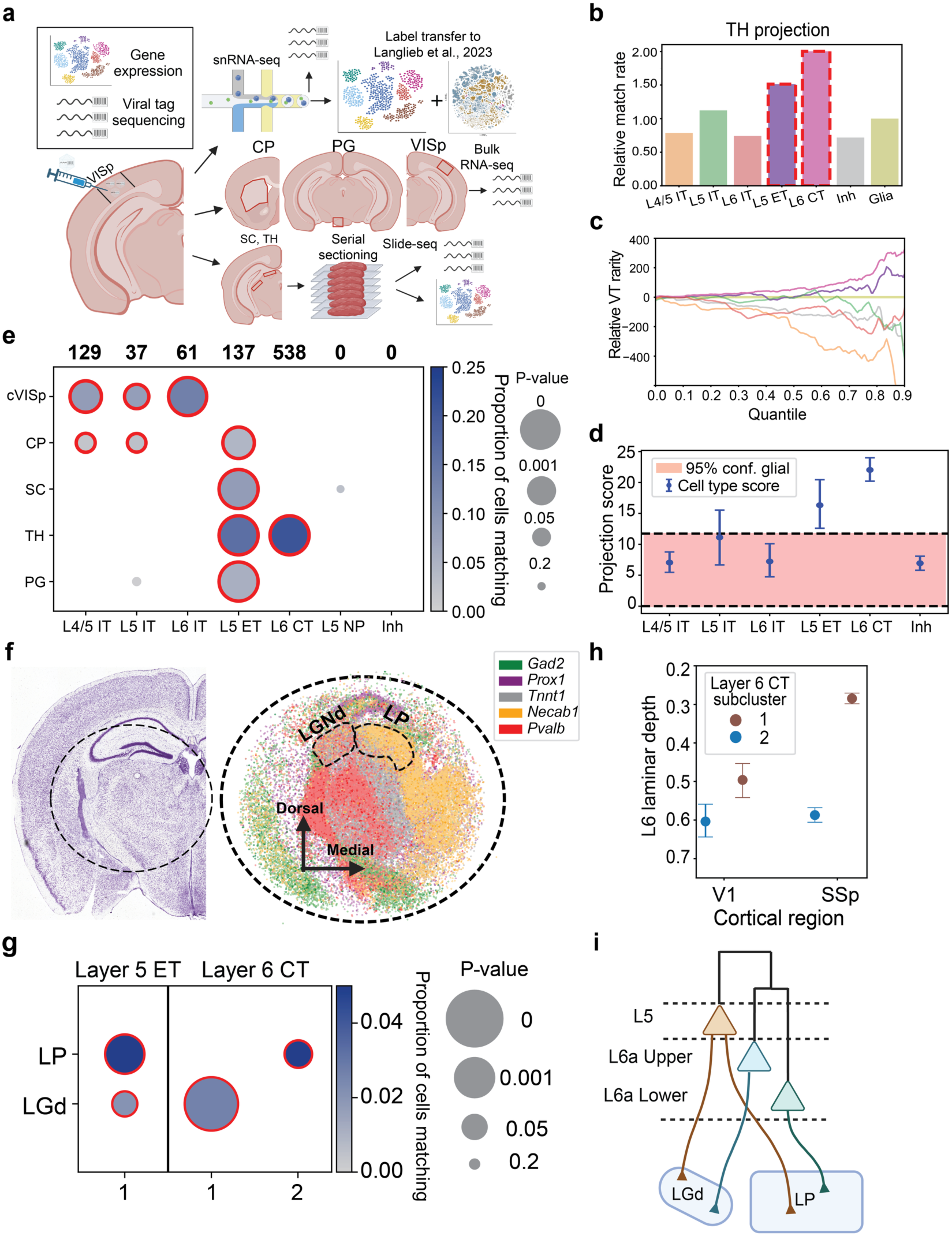
VISp projection mapping using Synapse-seq presynaptic barcoding. **a**, Experimental workflow and projection targets. Abbreviations: cVISp, contralateral VISp; CP, striatum; SC, superior colliculus; TH, thalamus; PG, pons. **b**, Barplot illustrating the VT matching rate to TH per cell type (scaled to glia). Red boxes: known thalamus-projecting types. **c**, ECDF of VT rarity in the AAV pool matching TH (normalized to glia). Positive values: VTs less common than in glia. **d**, Projection scores for excitatory neuron subclasses (whiskers: 2.5–97.5% quantiles; red shade: glial range). **e**, VISp subclass matching to major regions vs. glial null. Dot size is proportional to significance (1/p; red outline p < 0.05); color: matching proportion. Top: total matching cells. *n* (nuclei): L4/5 IT (1,342), L5 IT (361), L5 ET (509), L6 CT (2,597), L5 NP (256), Inh (2,044) (full statistical results reported in Supplementary Table 10). **f**, Thalamic nissl stain (left) and 15 aligned Slide-seq arrays showing LGd/LP boundaries (right). **g**, Projection matching of L5 ET and L6 CT clusters to thalamic nuclei. *n* (nuclei): ET (435), CT-1 (1,361), CT-2 (697). **h**, Laminar depth of L6 CT subpopulations in VISp and SSp (mean ± 95% confidence interval). **i**, Summary model: L5 ET targets both nuclei, while L6 CT subtypes show laminar and projection bias.

The two neuron classes known to project to thalamus–L5 ET and L6 CT–had the highest barcode matching rates in our thalamic Slide-seq dataset (Fig. 2b), and matched to comparatively rarer, more specific barcodes in the AAV pool (Fig. 2c). Leveraging this observation, we formulated a probabilistic model to distinguish true projections from technical noise. This framework penalizes overabundant barcodes that cause VT “collisions” (estimated at a rate of 13%) (Extended Data Fig. 4) and accounts for ambient RNA contamination based on empty-droplet profiles^33,34^. Crucially, we benchmarked significance against glial types as a statistical null (Methods), ensuring that projection calls exceed the false-positive baseline of non-projecting cells (Fig. 2d). Applied to all assayed targets, this conservative approach yielded significant matches for a total of 902 VISp cells, representing 16% of transduced excitatory neurons, and recovered the expected projection logic for each sufficiently transduced and captured cell type (Fig. 2e, Extended Data Fig. 3f and Methods). Specifically, L6 CT and L5 ET were the only thalamus-projecting classes, IT neurons projected exclusively to forebrain targets (contralateral VISp, striatum), and ET neurons projected across both forebrain and non-forebrain structures (Fig. 2e, Extended Data Fig. 5a). Consistent with prior work^35–36^, we observed that both L5 ET and L4/5 IT cells projected most strongly to the most dorsomedial part of the striatum (Extended Data Fig. 5b-f).

Finally, we assessed whether Slide-seq could resolve distinct cortical subtype connectivity to neighboring thalamic nuclei (Fig. 2f, Extended Data Fig. 6a,b). Indeed, a layer 5 ET subtype (Ex_Fezf2_Klhl1) projected to both LGd and lateral posterior nucleus (LP), while a superficial layer 6 CT subtype (Ex_Foxp2_Col5a1, “CT-1”) projected preferentially to LGd. Two transcriptomically related, deeper layer 6 CT types (Ex_Fezf2_Sebox and Ex_Fezf2_Cpa6, “CT-2”) projected preferentially to LP (jackknife p < 0.05) (Fig. 2g, Extended Data Fig. 6c-f). These findings mirror published reports of depth-dependent corticothalamic targeting by layer 6 neurons in the SSp^37,38^ (Fig. 2h), and suggest a general organizational rule: higher-order thalamic nuclei (e.g., LP) receive input from deeper L6a CT subtypes, whereas lower-order nuclei (e.g., LGd) are targeted by more superficial L6a CT subtypes (Fig. 2i). Broadly, our VISp circuit mapping demonstrates that presynaptic Synapse-seq can recover known projection motifs and resolve previously unknown fine topographies of molecularly defined cell types.

## Topographical mapping of anterior cortical collaterals to striatum and medulla

We next extended our mapping of long-range cortical circuitry to the anterior cortex^39^, characterizing cell type-resolved projections from this region to the striatum, thalamus, subthalamic nucleus, contralateral cortex, and medulla (Fig. 3a). From the injection site, we recovered 6,158 nuclei with joint RNA and VT profiles (mean 4.8 unique VTs per cell) (Extended Data Fig. 7). All significant projections, detected in a total of 689 of these cells (18% of all virally transduced excitatory neuron profiles) aligned with known projection anatomy^14,40^ (Fig. 3b, Extended Data Fig. 8a).

**Fig. 3:**
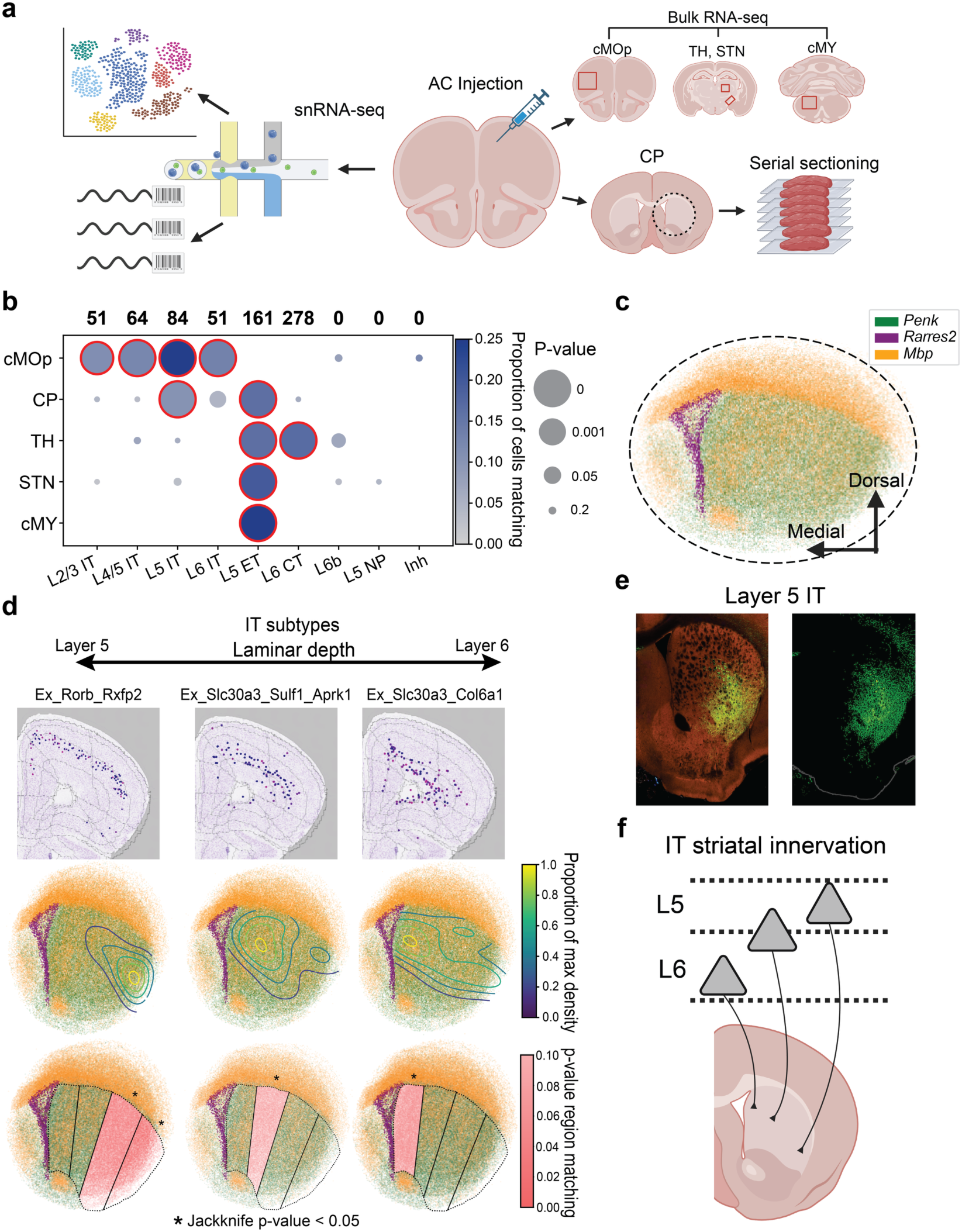
Systematic analysis of projections from the anterior cortex (AC) using presynaptic Synapse-seq. **a**, Schematic illustrating experimental workflow and surveyed dissection regions for AC injection experiment. cMOp, contralateral motor cortex; CP, striatum; TH, thalamus; STN, sub-thalamic nucleus; cMY, contralateral medulla. **b**, Dotplot depicting regional projection analysis between molecular subclass and each projection region. Red circles are drawn around significant projections (jackknife p-value < 0.05). Top: total matching cells per subclass. *n* (nuclei): L2/3 IT (458), L4/5 IT (513), L5 IT (289), L6 IT (388), L5 ET (308), L6 CT (1,648), L6b (87), L5 NP (160), Inh (1443) (full statistical results reported in Supplementary Table 10). **c**, Fourteen aligned striatal Slide-seq arrays plotted with spatially defined markers (Methods). **d**, Top, representative screenshots of laminar depth for each IT subtype matching to striatum^1^. Matching VTs from each IT subtype are shown as contour density maps (middle, Methods), and as regional heat maps of projection significance (bottom), where the red shading is scaled by jackknifed p-value (* p < 0.05). *n* (nuclei): Ex_Rorb_Rxfp2 (271), Ex_Slc30a3_Sulf1_Aprk1 (132), Ex_Slc30a3_Col6a1 (207) (Supplementary Table 10). **e**, Anterograde tracing experiments showing corticostriatal projections from MOs for a layer 5-specific Cre from the Allen Mouse Brain Connectivity Atlas (Experiment: 517313256). **f**, Proposed models of striatal innervation by IT cell type groups. IT types located more superficially project more lateral-ventrally to the striatum, while deeper laminar types project to the more medial-dorsal aspect of the striatum.

Leveraging Slide-seq data of the striatum (Fig. 3c, Extended Data Fig. 9), we found robust depth-dependent differences in the spatial distributions of projections from L5 compared with L6 IT cell types: the most superficial L5 IT subtype (Ex_Rorb_Rxfp2) targeted the ventrolateral striatum, the deepest L6 IT subtype (Ex_Slc30a3_Col6a1) innervated the dorsomedial region, and an intermediate subtype (Ex_Slc30a3_Sulf1_Aprk1) straddled the two (jackknife p-value < 0.05) (Fig. 3d, Extended Data Fig. 8b,c, Extended Data Fig. 9c and Methods). Sparse labeling of a layer 5-IT-specific Cre line independently confirmed this spatial specificity^40^ (Fig. 3e). Together, these data reveal a continuous relationship between cortical depth and striatal target position amongst IT corticostriatal neurons (Fig. 3f), an organizing principle difficult to capture in standard tracing experiments.

Prior work has indicated an additional medial-to-lateral topography in the anterior cortex in which ET neurons project differentially to multiple targets, including the striatum and medulla^39^, yet the exact transcriptomic identity of these projecting neurons remains unclear. By tracing copies of barcodes trafficked to multiple projection sites of a single neuron, Synapse-seq enables the detection of collateral projections. Subdividing our injection site into 12 medial-to-lateral dissections (Fig. 4a), we confirmed the reported organization of medial ET neurons projecting to the ventral medulla compared to more dorsally projecting lateral ET neurons (t-test difference in mean p < 0.001) (Fig. 4b). Furthermore, we observed a striking correlation in which neurons projecting to the ventral medulla sent collaterals to the dorsomedial striatum, while those with more dorsal medullary projections sent collaterals to ventrolateral striatum (Fig. 4c,d).

**Fig. 4:**
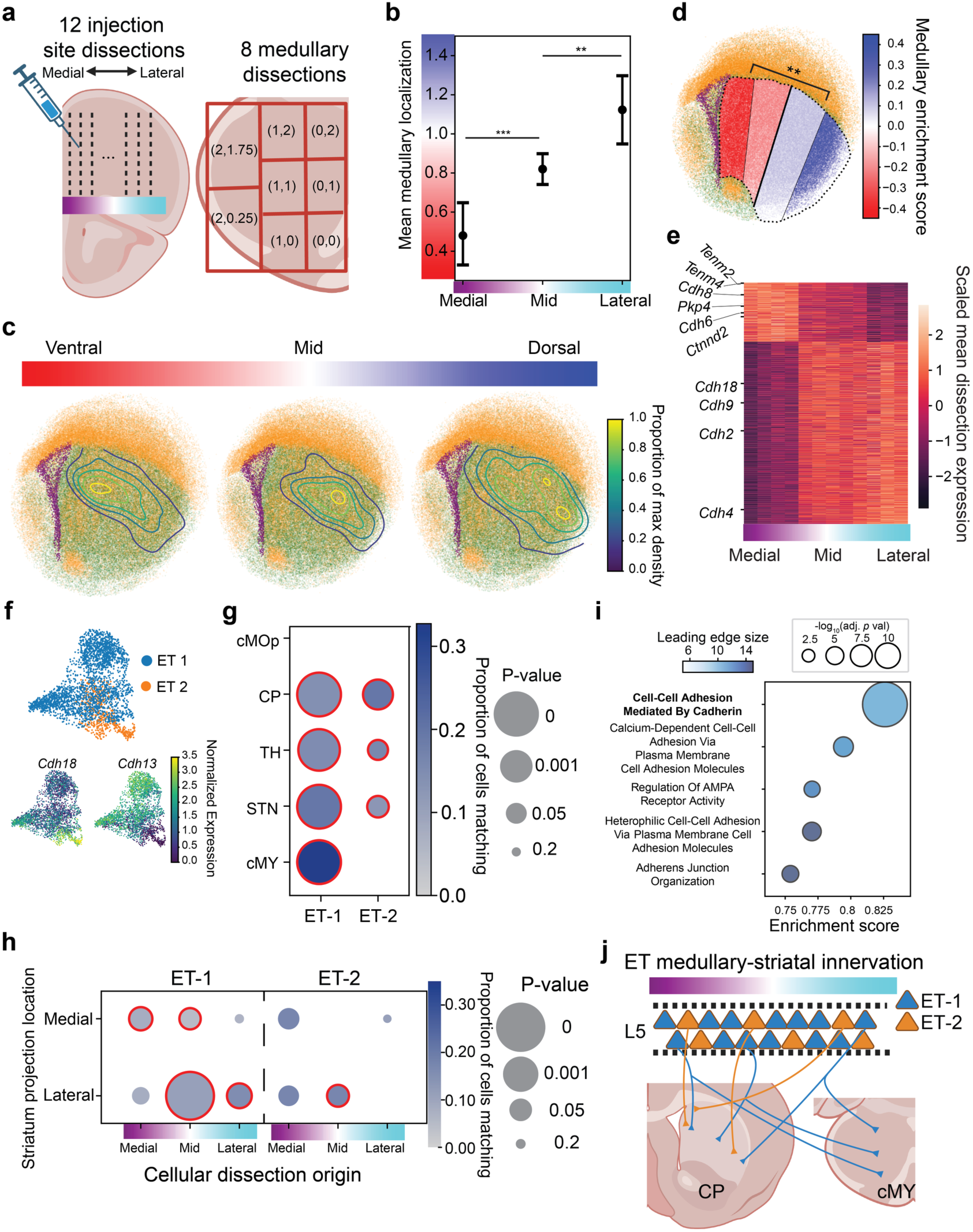
Medullary collaterals predict spatial innervation of L5 ET striatal projections. **a**, AC sampling strategy showing medial-to-lateral cortical dissections (left) and medullary dissections with assigned pseudocoordinates (right). **b**, Mean medullary projection location per cortical dissection group (mean ± 95% confidence interval). Permutation *t*-test: medial vs. mid (*t*=5.31, *p*<0.0001), mid vs. lateral (*t*=3.01, *p*<0.01). *n* (VTs): medial (52), mid (166), lateral (46). **c,** Striatal contour maps localizing VT matches to contralateral medulla, grouped by dorsal-ventral axis (Methods). **d**, Striatal enrichment of ventral (blue) vs. dorsal (red) medullary collaterals (** p < 0.01, difference in permutation test; *n* (VTs) = 331 medial, 256 lateral). **e,** Heatmap of genes correlating with the cortical medial-lateral axis (scaled log1p expression). Gradient indicates dissection location shown in **a**. Marked genes: *Tenm2*, *Tenm4,* and top GO term “Cell-Cell Junction Organization.” **f**, UMAP of ET subtypes (top) and *Cdh18*/*Cdh13* expression (bottom). **g**, Dotplots of projection matching of ET subtypes to target regions. *n* (nuclei): ET-1 (242), ET-2 (52) (full statistical results reported in Supplementary Table 10). **h,** Dotplot of projection matches by ET subtype, source dissection, and striatal projection regions defined in **d**. *n* (nuclei) for ET-1/ET-2: Medial (73/17), Mid (125/17), Lateral (44/18) (Supplementary Table 10). **i**, Top 5 GO terms from a GSEA analysis of differentially expressed genes between the two ET subclusters. **j**, Model of coordinated topography in medulla-projecting ET neurons.

Taking advantage of Synapse-seq’s ability to link gene expression to cortical positions and projections, we found that many genes varied systematically across this cortical axis, including synapse assembly genes such as *Tenm2* and *Tenm4*^41,42^ (Fig. 4e, Extended Data Fig. 10a-c, Supplementary Tables 1,2). Furthermore, we identified two transcriptomically diverse ET groups, one of which sends collaterals to the medulla (“ET-1”) and another that does not (“ET-2”) (Fig. 4f,g, Extended Data Fig. 10d,e). Only the medulla-projecting ET-1s showed strong coupling between cortical medial-to-lateral position and striatal projection location (Pearson R 0.57, p-value < 0.01) (Fig. 4h, Extended Data Fig. 10f). Differential expression identified prominent variation in cell-adhesion gene signatures (Fig. 4i, Supplementary Tables 3-4), with *Cdh13* and *Cdh18* exhibiting particularly selective expression (Fig. 4f). Thus, cadherin-defined ET subtypes are characterized by selective medullary innervation and separable striatal topographies (Fig. 4j). These analyses demonstrate that orthogonal spatial dimensions–laminar depth for IT neurons versus medial-lateral position for ET neurons–cooperate with specific transcriptional identities to structure the output of the anterior cortex.

## Postsynaptic barcoding recapitulates cell type-specific features of dendritic morphology

Dendritic integration is also critical to neural circuit function, yet methods for mapping these input sites are often low-throughput or lacking in precise cell type identification. To characterize postsynaptic neuroanatomy with Synapse-seq, we injected the PSD95.FingR-guided mRNA barcoding system into the hippocampus (Fig. 5a), a tissue with functionally-relevant dendritic architecture^43^. In an aligned Slide-seq dataset, we sampled gene expression and VTs from each hippocampus subfield (CA1, CA2, CA3, DG) (Extended Data Fig. 11a-c, Supplementary Table 5 and Methods). Compared to a randomly permuted distribution, we observed strong spatial colocalization of VT matches, consistent with barcode labeling of individual cells (Extended Data Fig. 11d). After correcting for VT collisions (estimated at a rate of 14.6%), we reconstructed 18,742 distinct “VT trees” representing approximately 7,967 pyramidal and 1,897 dentate granule cells (Fig. 5b-c, Extended Data Fig. 11e and Methods).

**Fig. 5:**
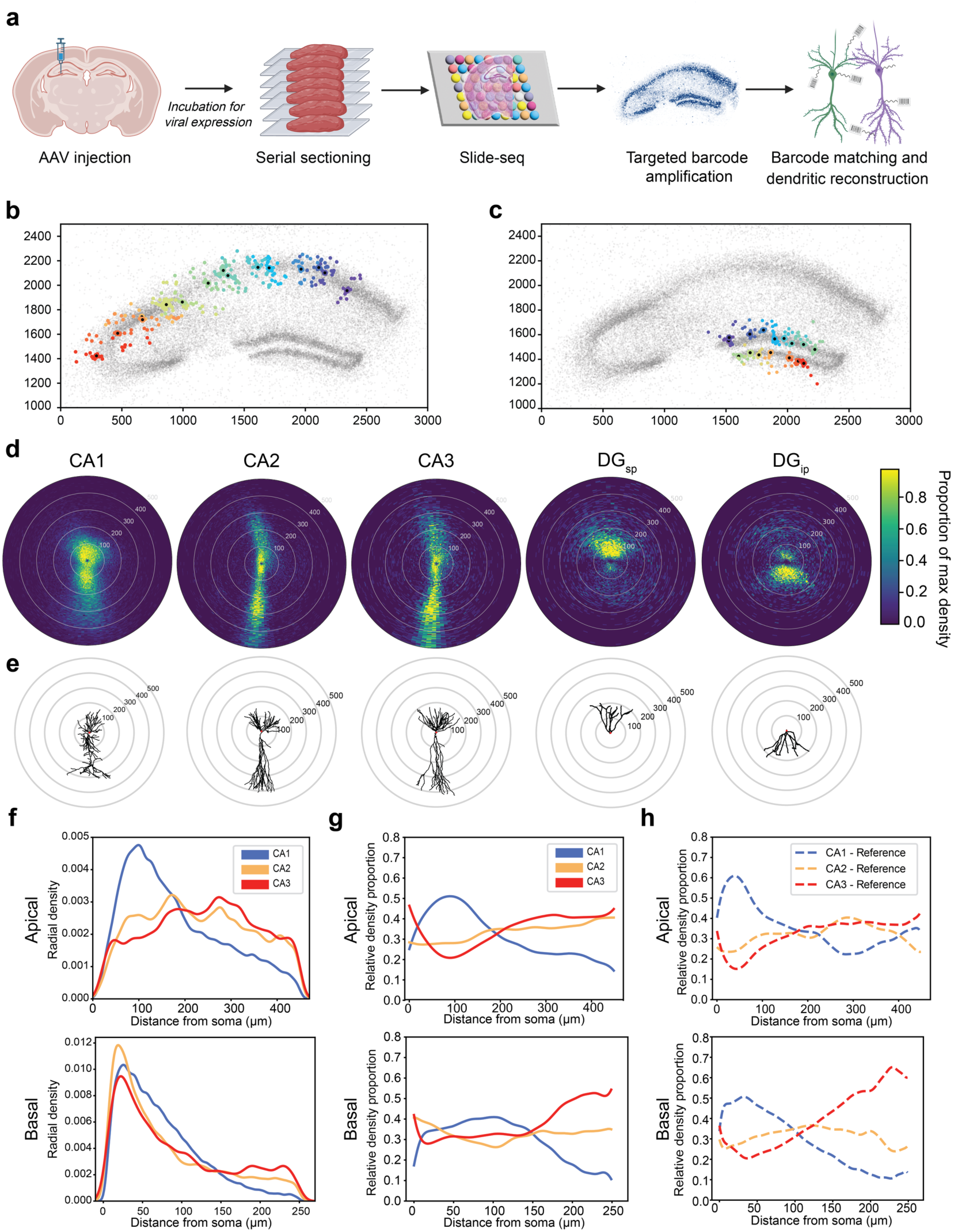
Postsynaptic Synapse-seq barcoding reveals distinct dendritic architectures of hippocampal pyramidal and dentate granule neurons. **a**, Experimental schematic of hippocampal postsynaptic barcoding approach. **b-c**, Example subset of DBSCAN-clustered VT trees across CA, **b**, and DG, **c**, subfields, showing estimated soma placement (black dot) and synaptic matches (colored dots) against background of all Slide-seq beads (grey dots, intensity reflects total number VT matches per bead). Axis distances in the coronal plane are shown in μm; anterior-posterior z-dimension is collapsed for visualization purposes; unique VT trees are colored according to their position along the soma layer. **d**, Polar coordinate plots (*r*, *ϕ*), depicting consensus dendritic morphology of the CA1, CA2, CA3, suprapyramidal DG (DG_sp_), and infrapyramidal DG (DG_ip_) (Methods). Each bin is colored based on the proportion of total density of VT matches contained. **e**, Representative reference dendritic reconstructions^46^ from the same region defined directly above in **d**. **f**, Radial density analysis comparing the distribution of three-dimensional radial distance from the soma of VT matches for the apical (top) and basal (bottom) layers respectively, across pyramidal cell types. **g**, Relative normalized density proportions of CA1, CA2, and CA3 pyramidal neurons, in their apical (top) and basal (bottom) layers (Methods). **h**, Relative normalized density plots, processed as in **g**, of the reconstructed reference dataset^46^.

Aggregating VT trees onto shared polar coordinate axes (Methods) recapitulated stereotyped morphologies, including the apical-basal asymmetry of pyramidal neurons, and the directionally opposed, “Y-shaped” dendrites of suprapyramidal and infrapyramidal dentate granule cells^44,45^ (Fig. 5d, Extended Data Fig. 11f-h). To quantify these barcoded postsynaptic distributions, we measured the radial distance of VT matches from their somas and compared our results to microscopy-based reconstructions^46^ (Fig. 5e-h and Methods). Synapse-seq accurately resolved fine-grained morphological features, including the distinctive proximal apical density of CA1, and the progressive anterior-to-posterior shift in dendritic orientation from CA1 to CA3 (Fig. 5f-h, Extended Data Fig. 11i,j). Our postsynaptic barcoding also recovered a previously reported distal bias in the dendritic distributions of suprapyramidal dentate granule cells, relative to their infrapyramidal counterparts^47^ (Extended Data Fig. 11k).

Although superficial and deep hippocampal pyramidal neurons exhibit distinct genetic^48^, morphological^49^, and physiological^50^ properties, coordinated multimodal analysis across subfields remains technically challenging. Exploring this in our data, we first identified differentially expressed genes along the superficial-deep axis in CA1 and CA3, but not CA2 (Fig. 6a, Supplementary Table 6). These genes were highly concordant with published datasets^1,48,51,52^ and significantly enriched for synaptic gene ontology terms; notably, postsynaptic terms were significant only in CA1 (Fig. 6b, Extended Data Fig. 12a, Supplementary Table 7).

**Fig. 6:**
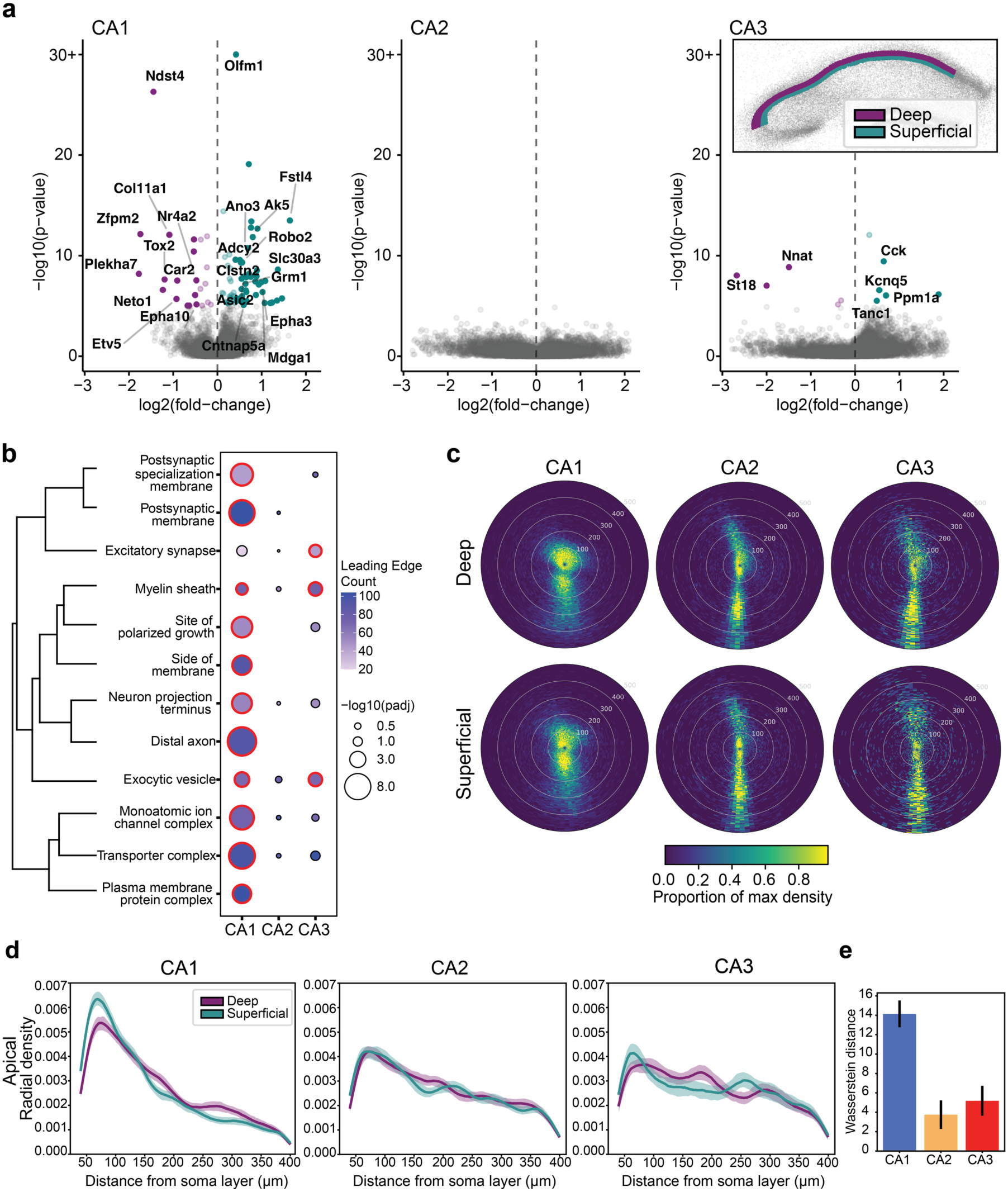
Barcoding-based ascertainment of transcriptomic and morphological variation along the superficial-deep axis. **a**, Volcano plots depicting differential expression (DE) results, across CA1 (left), CA2 (middle), and CA3 (right). Points are colored based on their enrichment in the superficial or deep layer (inset), with significant (Bonferroni-adjusted p-value <0.05) DE genes labeled in bold terms indicating concordant differential expression in reference datasets^1,48^ (Methods)**. b**, GSEA-based GO cellular compartment term enrichment of superficial-deep differential expression results in each hippocampal subfield. Dendrogram (left) denotes hierarchical clustering of GO terms by semantic similarity^63^. **c**, Polar coordinate plots of VT match density as in Fig. 5d, separated by their soma positions in the deep (top) or superficial (bottom) layer across hippocampal subfields. **d**, Radial density analysis (Methods) of the apical layer, comparing distance distributions between superficial and deep cells, across pyramidal neuron types. Distances reflect Euclidean distance between a VT match and the intersection point at which the VT tree crosses the soma layer boundary. 95% confidence intervals shaded. **e**, Wasserstein distances computed between the apical distributions of superficial and deep VT density distributions, across CA subfields. Error bars reflect standard error computed by bootstrapping (Methods).

Morphologically, deep-layer neurons across subfields shared common features, including a higher ratio of basal-to-apical VTs (51-56% vs. 32-37%), and a selective reduction in proximal apical VT density that corresponds to the characteristic unbranched initial segment, or “neck”, of deep pyramidal dendrites^53,54^ (Fig. 6c, Extended Data Fig. 12b-d). Comparing apical radial density distributions beyond the soma layer revealed that superficial CA1 neurons possess a uniquely high proximal VT density relative to deep cells, whereas superficial-deep differences in CA2 and CA3 were more subtle (Fig. 6d,e, Extended Data Fig. 12e and Methods). These results mirror our transcriptomic findings, identifying CA1 as the subfield with the most pronounced superficial-deep heterogeneity, and establishing Synapse-seq as a platform for coupling dendritic architecture to transcriptional identity at scale.

## Discussion

Neurons can be classified by many modalities, including gene expression^4,13,14^, spatial localization^55^, and projection logic^7^. By profiling transcriptomes and synaptic anatomy from the same neurons at scale, Synapse-seq provides a tractable methodology for establishing integrative taxonomies. Using our system, we successfully mapped fine-grained projection targets and dendritic morphologies to molecularly-defined neurons.

Our application of Synapse-seq revealed circuit organizations that are invisible to conventional tracing approaches. In VISp, we resolved layer 6 CT cells into two molecularly distinct sublayers projecting to different thalamic nuclei. This suggests a conserved logic where superficial layer 6 subtypes project to first-order nuclei (e.g. LGd), while deeper subtypes target higher-order nuclei (LP). Similarly, in the anterior cortex, we identified cadherin-defined subtypes of L5 ET neurons that segregate by their collateral projections to the medulla, and L5 IT neurons with striatal projections that segregate by their cortical sublayer of origin. These findings demonstrate that orthogonal spatial dimensions–laminar depth and medial-lateral position–differentially organize the outputs of distinct cortical projection classes.

Applying our subcellular barcoding approach to the postsynaptic compartment, we resolved the three-dimensional apical and basal dendritic arbors of molecularly-defined excitatory hippocampal cell types. Relative to other hippocampal subfields, the position of CA1 somas along the superficial-deep axis showed the strongest association with postsynaptic gene signatures and morphological heterogeneity. Beyond the present dataset, our technology enables future comprehensive profiling of other key spatial dimensions–including the proximal-distal^56^ and dorsal-ventral^57^ axes. Finally, future integration of single-cell-resolution spatial tools like Slide-tags^48^ will enable even more precise dissection of intermixed cell types, and application to regions like the cortex that lack the spatial segregation of hippocampal cell bodies and synaptic layers.

Beyond these immediate findings, Synapse-seq offers a versatile platform for functional genomics. The system’s modularity enables barcoding of other subcellular compartments, such as mitochondria, axon initial segments, or primary cilia. Coupling these compartment-specific tags with single-cell transcriptomics and delivery of Perturb-seq–style guide-RNA libraries^58^ should enable pooled CRISPR screens on subcellular readouts such as presynaptic-terminal density or spatial distribution. Identifying genetic regulators of axon guidance and synaptic arborization using this approach would allow for the systematic dissection of neural circuit assembly *in vivo*.

While Synapse-seq has numerous direct uses, future improvements and modifications will further extend its reach. First, while we tested Synapse-seq’s ability to robustly and specifically label synaptic distributions of several cell types across major brain areas, its performance in other brain areas, such as hindbrain, remains to be established. Second, in our presynaptic experiments, we used probabilistic filtering to suppress barcode collision noise; consequently, sparse projections may fall below the current detection threshold, an issue that should improve with larger, more diverse AAV barcode libraries. Finally, our spatial barcoding achieves the 10-micron resolution specified by the bead diameter of Slide-seq arrays^31^; by altering the barcode architecture to permit hybridization-based barcode detection, Synapse-seq can be paired with high-resolution, microscopy-based spatial transcriptomics to resolve fine-scale features like individual dendritic spines.

Ultimately, a primary advantage of Synapse-seq is scalability. By leveraging AAVs and high-throughput sequencing, the same workflow can be applied across various stages of development, disease models, and species. Recent advances in systemic AAV engineering^59–62^ will facilitate the delivery of these constructs to the entire brain, enabling the generation of whole-brain maps across multiple individuals. The resulting paired datasets of transcriptomic identity and synaptic architecture will provide a quantitative scaffold for comparative neurobiology and for pinpointing the specific cells and synapses most vulnerable to aging and disease.

## Resource Availability

Processed data is publicly available in a Google Cloud Storage bucket: https://console.cloud.google.com/storage/browser/macosko_public/Synapseseq_data_2025. Custom code and vignettes for both presynaptic and postsynaptic analyses can be accessed at: https://github.com/MacoskoLab/SynapseSeq_Analysis. Plasmids will be available by time of publication on Addgene: https://www.addgene.org/Evan_Macosko/.

## Acknowledgments

We thank Gord Fishell, Evan Feinberg, and Fei Chen for helpful discussions. We thank Matthew Shabet, Mukund Raj, and all members of the Macosko laboratory for helpful discussions and feedback. We thank Pam Brauer, John Harvey, and Alexander Svanbergsson for help with virus production. This work was supported by the Life Sciences Research Foundation (to M-JD), the Merkin Fellowship at the Broad Institute (to EZM), the CONNECTS program within the NIH BRAIN Initiative (via grants RF1MH130468 and U01NS136398 to EZM), and the Stanley Center for Psychiatric Research. BZ and KC were supported by Brain Initiative awards funded through the National Institute of Mental Health (UH3MH120096 and UF1MH130701 to BED), the NIH Common Fund and the National Institute of Neurological Disorders and Stroke through the Somatic Cell Genome Engineering Consortium (UG3NS111689 to BED), the National Institute of General Medical Sciences (T32GM007753 and T32GM144273 to AU), and the Stanley Center for Psychiatric Research.

## Author Contributions

Study conception and design: M-JD, DA, BS, BED, EZM. Performed experiments: AU, M-JD, MK, JP, AB, SG, JL, NN, KB. Performed analysis: AU, JS, MK. Design and production of barcoded beads: VK. Design and production of barcoded AAV: M-JD, BZ, KC, BED. Provided reagents: DA. M-JD, JS, AU, and EZM wrote the manuscript with contributions from all co-authors.

---

**Extended Data Fig. 1:**
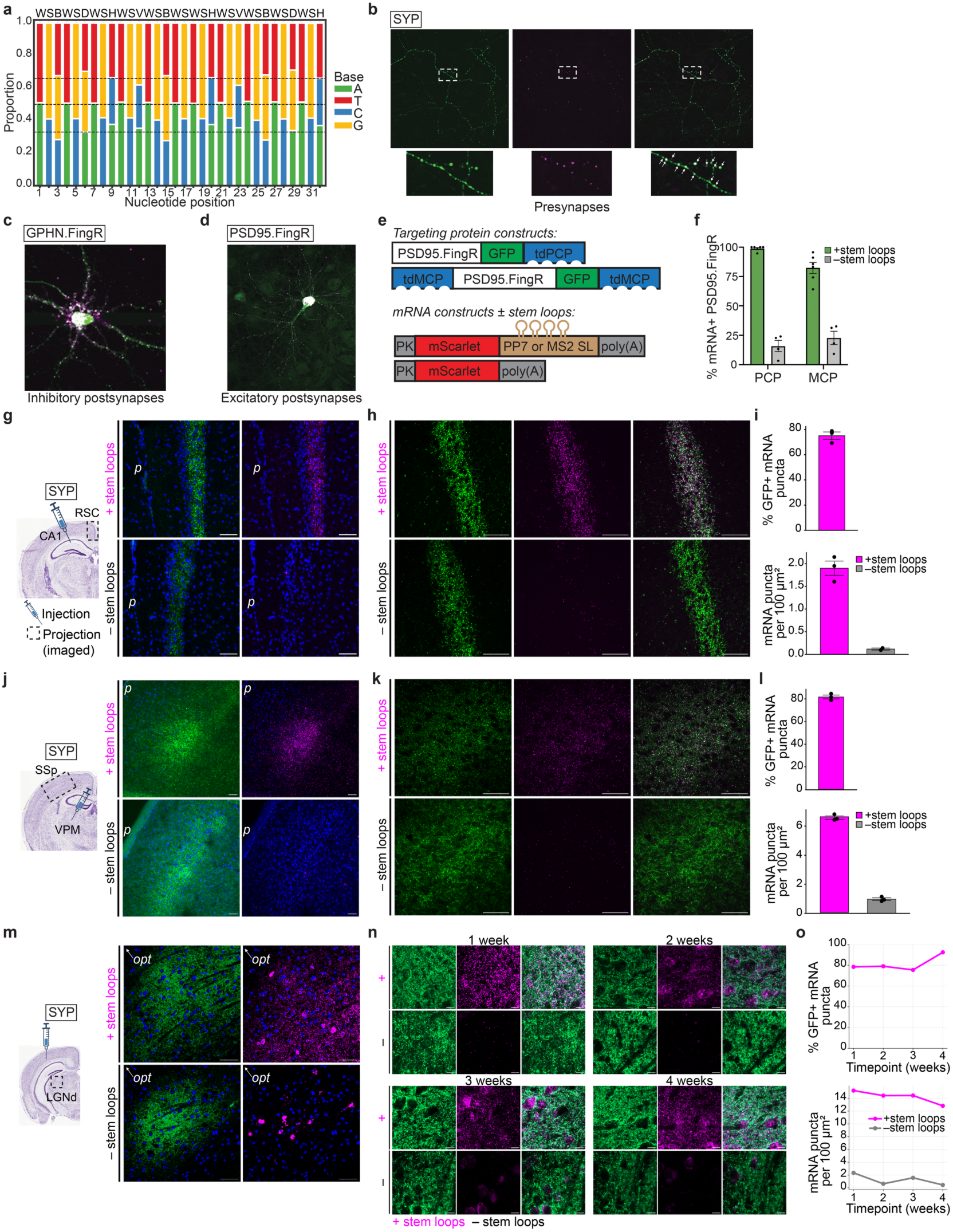
In-depth characterization of optimized Synapse-seq technology for labeling subcellular compartments across circuits and over time. **a**, Barcode nucleotide balance across the AAV library (dotted lines: 0.67, 0.5, 0.33). **b-f**, *In vitro* validation. Targeting protein (green, GFP) and *mScarlet* mRNA (magenta, HCR smFISH) in neurites (**b**) and dendrites (**c**,**d**). Inset in **b**: high-magnification presynapses. **e**, Constructs for comparing tdPCP-PP7 vs. tdMCP-MS2 targeting and stem loop dependence. **f**, Trafficking efficiency (% GFP with colocalized mRNA puncta) with (green, n=6) or without (grey, n=4) stem loops. **g**-**o**, *In vivo* characterization of hippocampo-retrosplenial (**g**-**i**), thalamocortical (**j**-**l**), and corticothalamic (**m**-**o**) projections. Images show SYP protein (green, GFP) and mRNA (magneta, RNAscope smFISH) in the RSC (**g**,**h**), SSp (**j**,**k**), and LGd (**m**,**n**) at 20x (scale bars 50 μm) and 63x (scale bars 10 μm) with (top) or without (bottom) stem loops. **i**,**l**, Quantification of labeling specificity and density (mean ± s.e.m.; n=2-3 per condition as shown). **n**,**o**, Stability in LGd over 1-4 weeks of viral incubation (anti-GFP IHC in **n**), quantified as in **i** (**o**; 1 week (n=3 per condition), 2-4 weeks (n=1 per condition). Cell body labeling in LGd (consistent with AAV9 tropism) was excluded from quantification.

**Extended Data Fig. 2:**
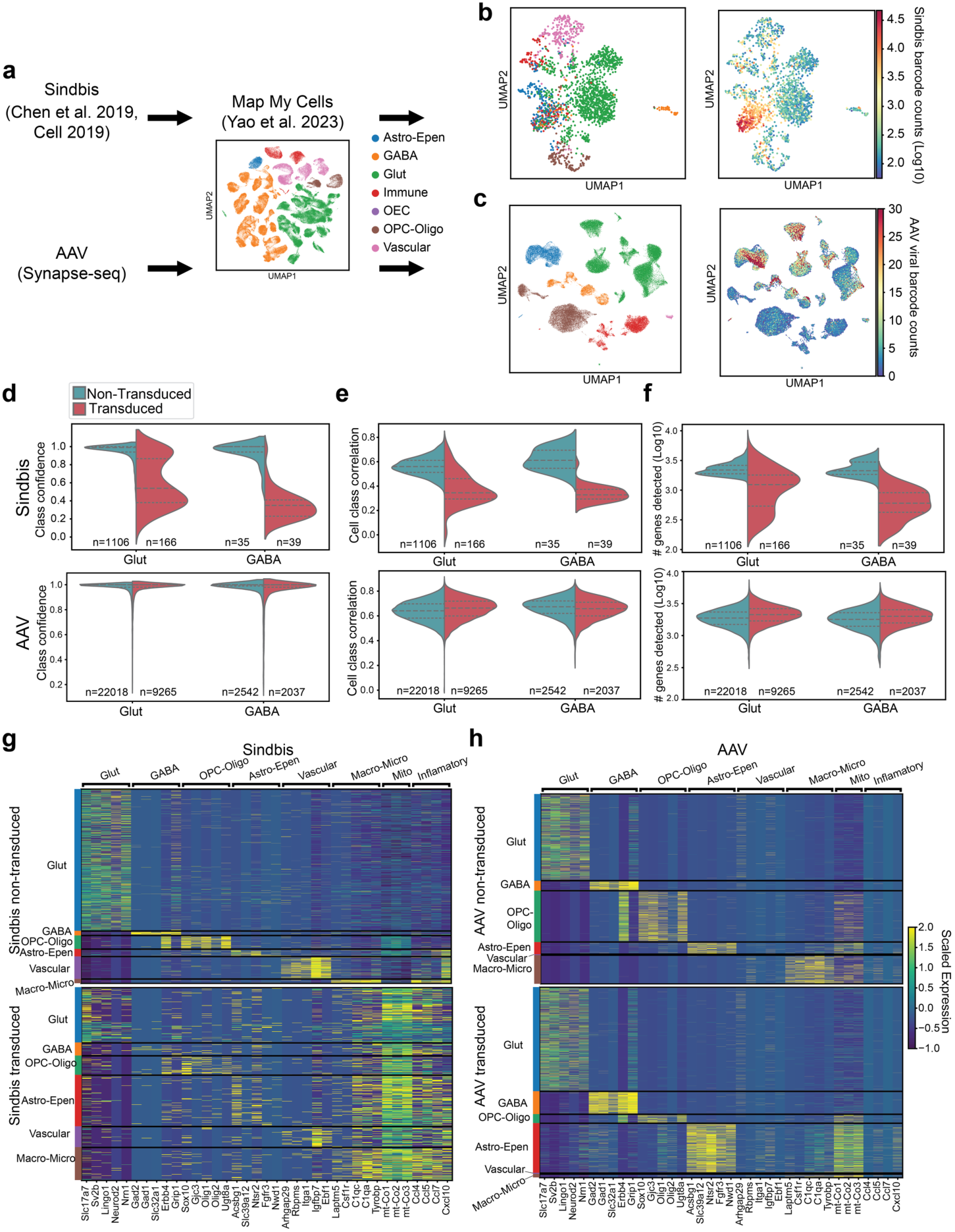
Transcriptomic impact of AAV-based mRNA barcode delivery in cortex. **a**, Schematic of label transfer from an annotated reference onto AAV- and Sindbis-transduced datasets. **b**, UMAP of Sindbis-transduced cells, colored by predicted cell class (left) and log10 number of viral barcodes per cell (right). **c**, UMAP of AAV-transduced cells, colored by predicted cell class (left) and number of viral barcodes per cell (right). **d-f**, Split violin plots comparing non-transduced (left) versus transduced (right) cells in both Sindbis (top row) and AAV (bottom row). Plots compare cell-type prediction confidence (**d**) correlation to the reference cell-class profile (**e**), and number of genes detected per cell (**f**). **g-h**, Heatmaps of scaled expression of cell-class markers, mitochondrial genes, and inflammatory signaling markers (Methods). Cells are grouped by predicted cell class for non-transduced (top) versus transduced (bottom) in both AAV (**g**) and Sindbis (**h**) datasets.

**Extended Data Fig. 3:**
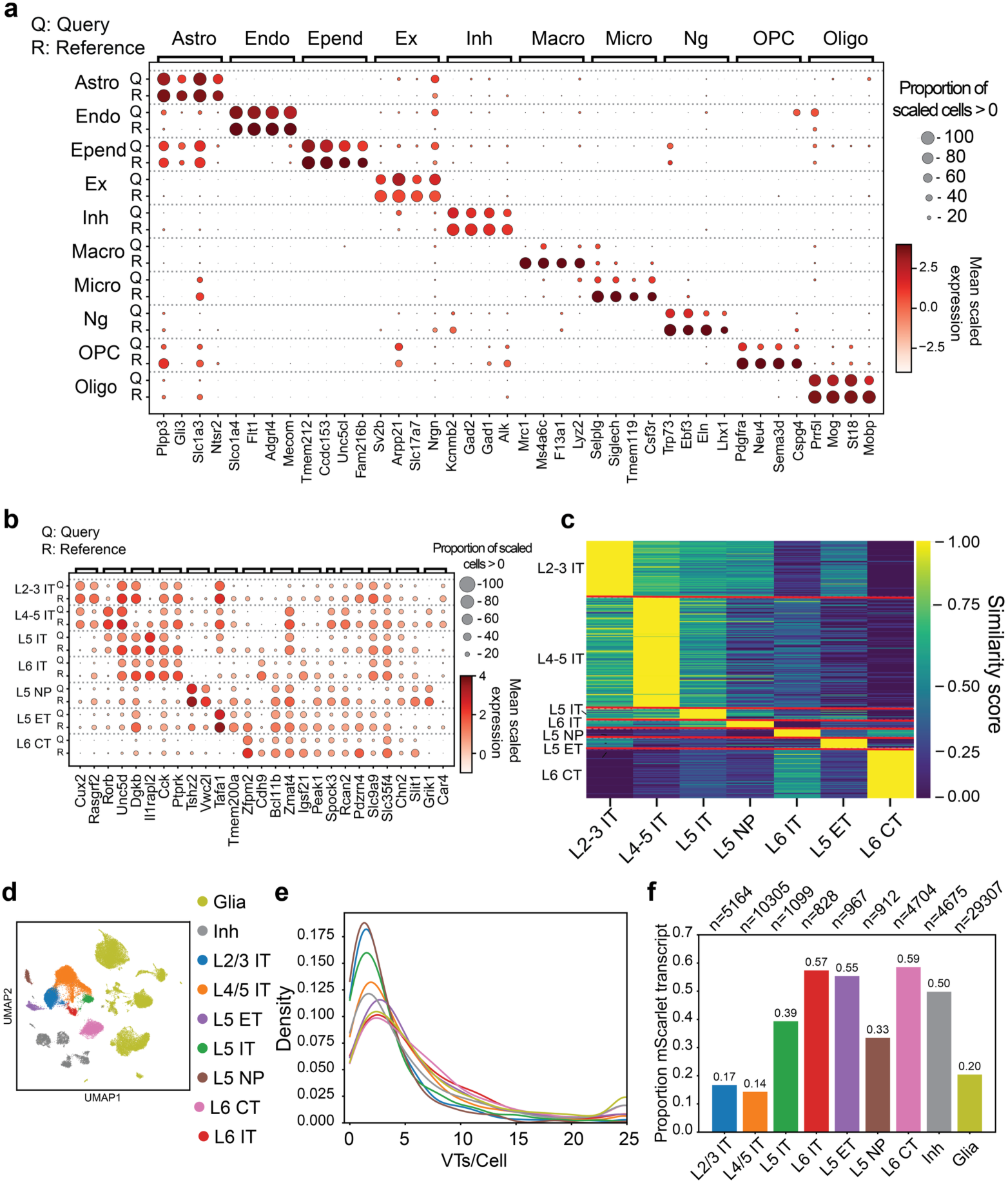
Cell type identification and viral barcode expression of sampled VISp cells. **a-b**, Dotplots depicting expression of cell type markers for cells from VISp injection experiment. Markers of cell class (**a**), and markers of excitatory neuron subsets (**b**), were computed on the reference dataset after sampling to match the predicted cell type proportions of the query dataset (Methods). Astrocyte (Astro), Endothelial (Endo), Ependymal (Epend), Excitatory neurons (Ex), Inhibitory neurons (Inh), Macrophages (Macro), Microglia (Micro), Neurogenesis (Ng), Oligodendrocyte progenitor (OPC), Oligodendrocytes (Oligo). **c**, Heatmap displaying gene expression similarity (Methods) for each predicted query cell to each reference subclass. **d**, UMAP embeddings of snRNA-seq cells, colored by type. **e**, Distributions of VTs per cell for each neuronal subclass, colored according to **d**. **f**, Barplot of the proportion of cells positive for an mScarlet viral transcript. The total number of cells in each subclass is noted above each column.

**Extended Data Fig. 4:**
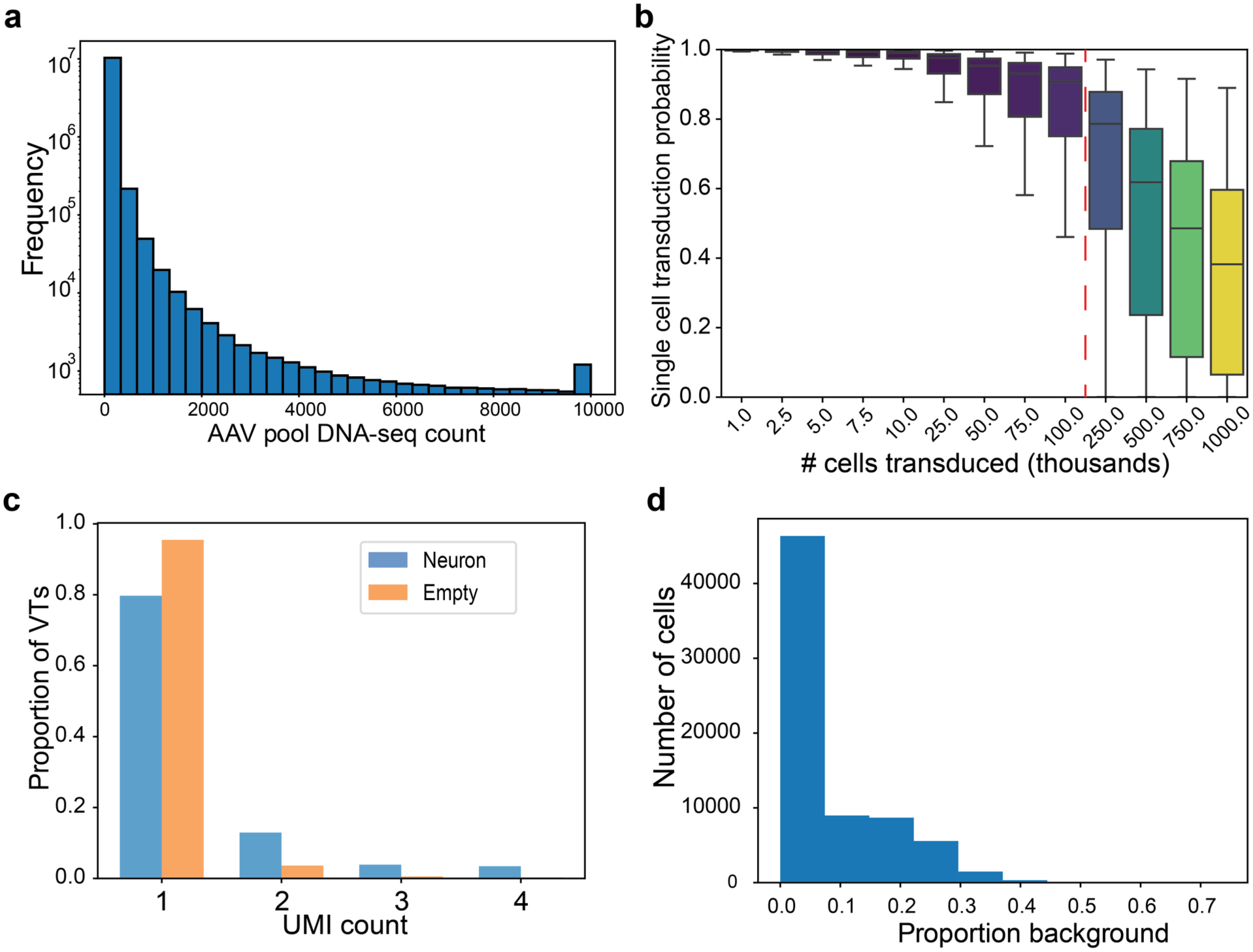
Analyses of VT barcode representation in the viral pool and snRNA-seq assays. **a**, Histogram depicting distribution of individual VT barcode counts detected in the AAV DNA-seq pool. **b**, Model of probability distributions that a given VT in the pool will transduce a single cell (versus multiple cells), computed at varying numbers of total cellular transduction events (x-axis). Whiskers indicate the 0.025-0.975 quantiles for each simulation (Methods). The dashed red line indicates the estimated total number of cells transduced (181,000) in the VISp experiment. **c**, Barplot showing the proportion of VTs in either *mScarlet* positive neurons or empty droplets with a certain UMI count. **d**, Distribution of the proportion of molecules attributable to background, estimated by Cellbender (Methods).

**Extended Data Fig. 5:**
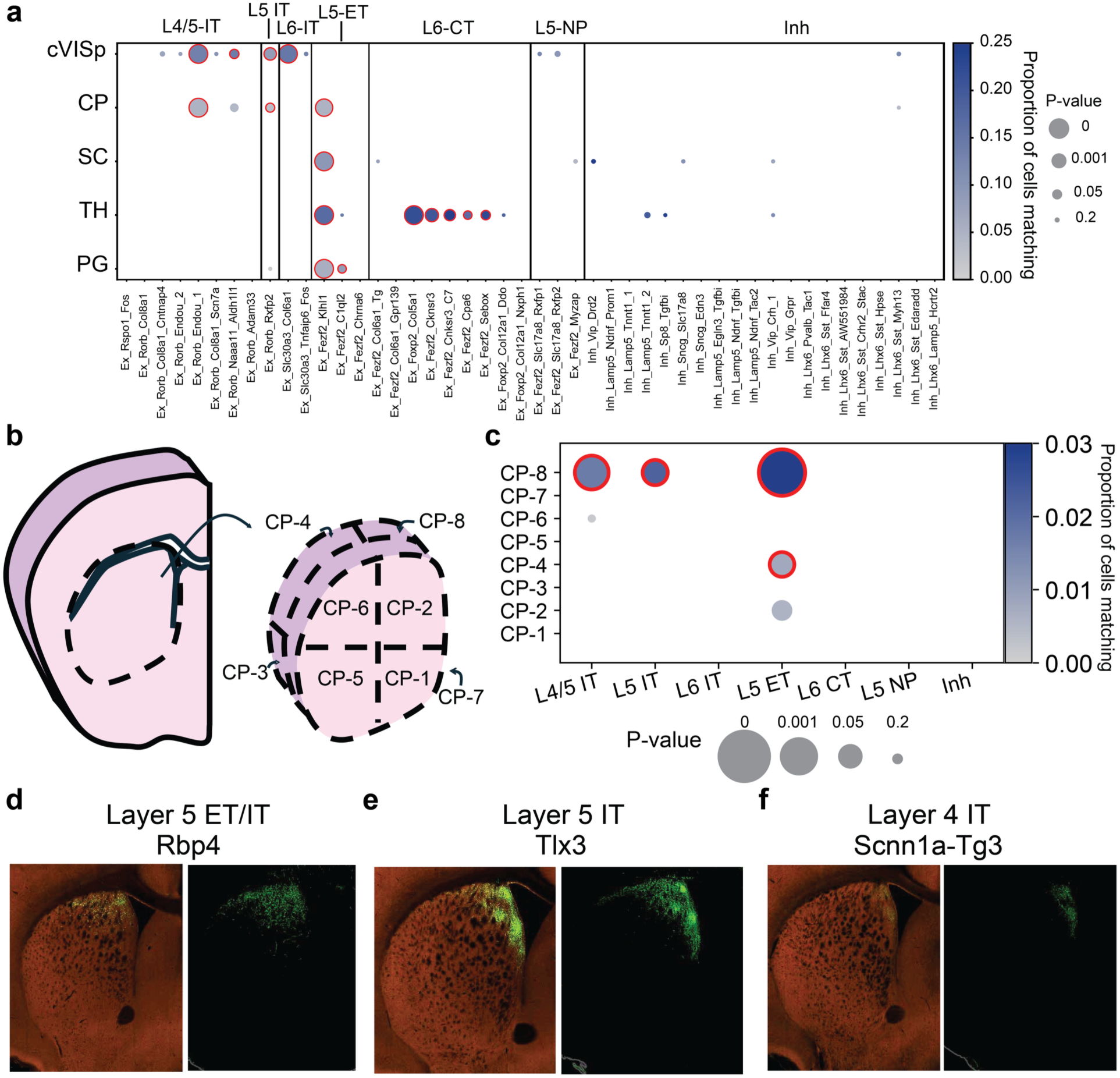
Additional analyses and validations of VISp projections measured by presynaptic Synapse-seq. **a**, Projection matching for VISp clusters, grouped by subclass (full statistical results reported in Supplementary Table 10). cVISp, contralateral VISp; CP, caudoputamen; SC, superior colliculus; TH, thalamus; PG, pons. **b**, CP dissection diagram. **c**, Subclass projection matching to subdissections in **b** (Supplementary Table 10). **d**-**f**, Allen Connectivity Atlas traces of VISp-striatal projections from L5 (*Rbp4-Cre*, d; ID: 65663288), L5 IT (*Tlx3-Cre*, e; ID: 529692491), and L4 IT (*Scnn1a-Tg3*, f; ID: 166324604) neurons. Panel **f** was vertically reflected for consistency.

**Extended Data Fig. 6:**
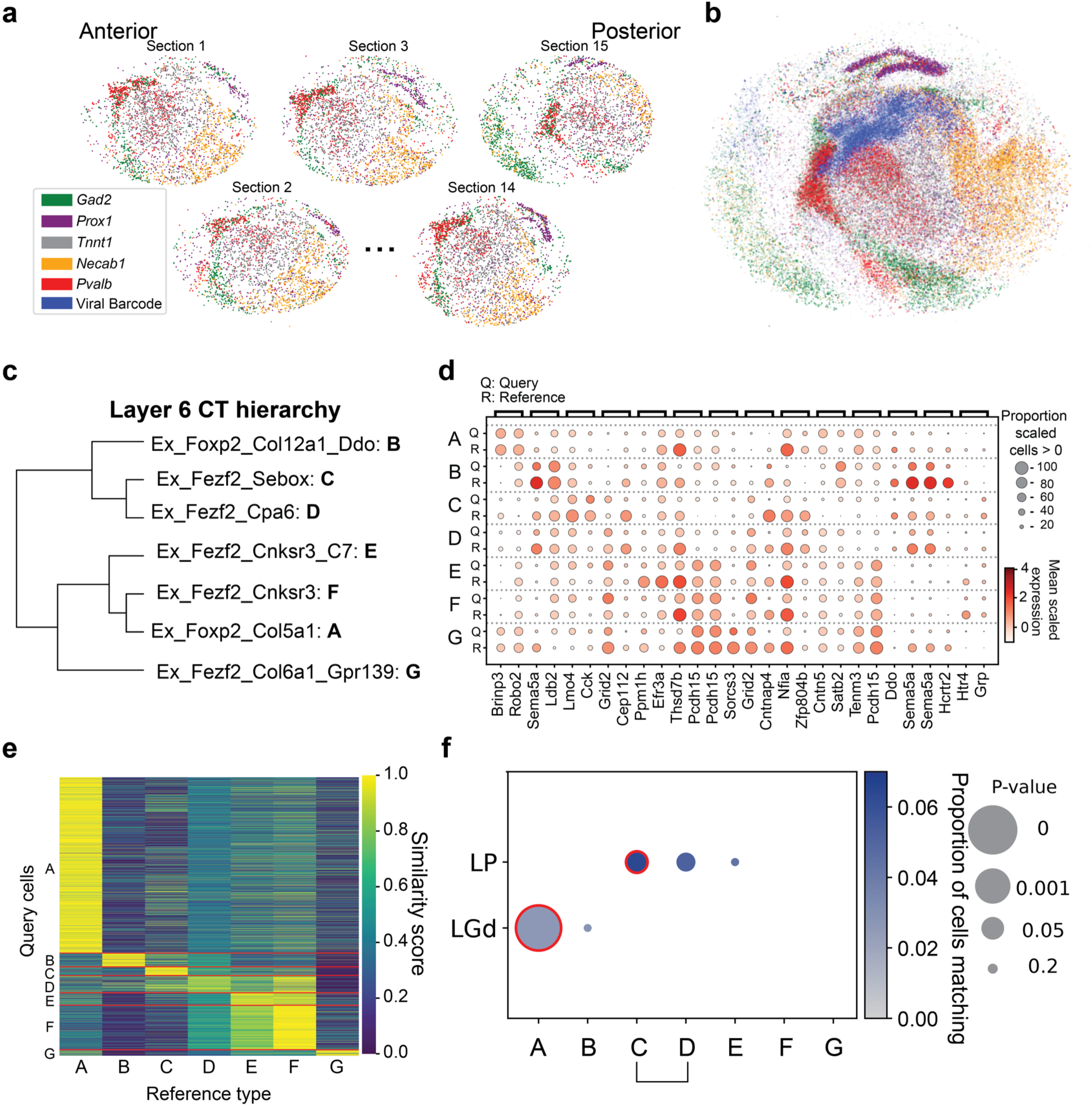
Thalamic projections of L6 CT subtypes. **a**,**b**, Slide-seq arrays arranged anterior-posterior (**a**) and collapsed along the z-axis (**b**), colored by spatial marker genes (left). **c**, Dendrogram of Layer 6 CT clusters from the whole-brain atlas^1^; letter labels are used in subsequent panels. **d**,**e**, Cluster identity validation via dotplot (**d**) and similarity heatmap (**e**). **f**, Projection matching of L6 CT subtypes to the LP and LGd (full statistical results reported in Supplementary Table 10). Types C and D were merged into CT-2 based on the dendrogram in **c**. LGd, dorsal lateral geniculate nucleus; LP, lateral posterior.

**Extended Data Fig. 7:**
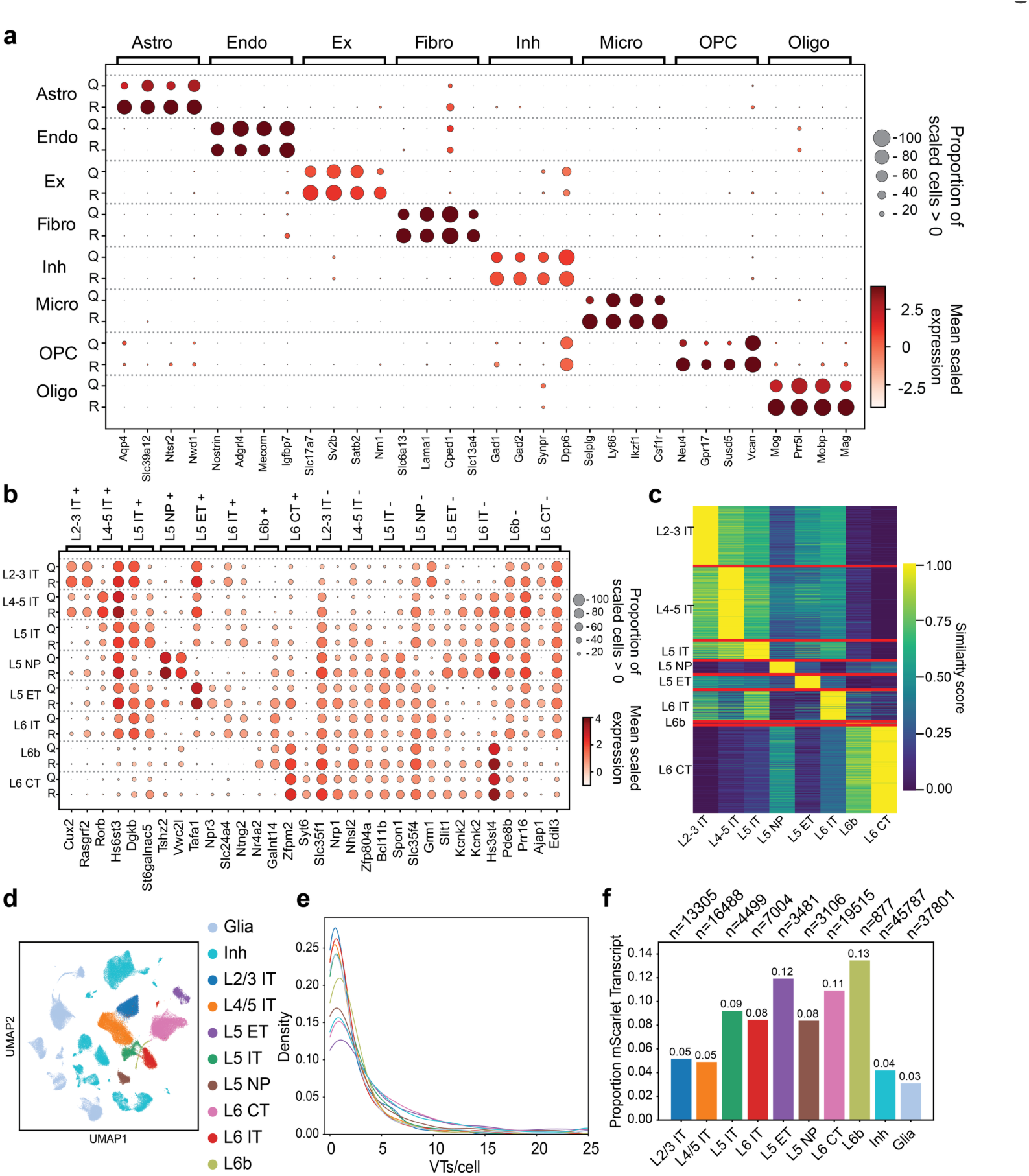
Cell type and viral barcode expression of sampled AC cells in presynaptic Synapse-seq experiment. **a**-**b**, Dotplots showing the correspondence of molecular markers (Methods) between the predicted cell classes (**a**) and subclasses (**b**) for cells dissociated from the AC. Astro, Astrocyte; Endo, Endothelial; Epend, Ependymal; Ex, Excitatory neurons; Inh, Inhibitory neurons; Macro, Macrophages; Micro, Microglia; NG, Neurogenesis; OPC, Oligodendrocyte progenitor; Oligo, Oligodendrocytes. **c**, Heatmap showing similarity score based upon scaled correlations (Methods) between each query excitatory neuron and possible subclasses, using reference-computed markers for each subclass (Methods). **d**, UMAP embeddings of snRNA-seq profiles from AC, colored by class (glia, inhibitory) or subclass (excitatory neurons). **e**, Density plot displaying distributions of number of VTs detected per cell, colored by legend in **d**. **f**, Barplot showing rate of *mScarlet* transcript positivity for each cell, in each subclass. Top: total number of cells per type.

**Extended Data Fig. 8:**
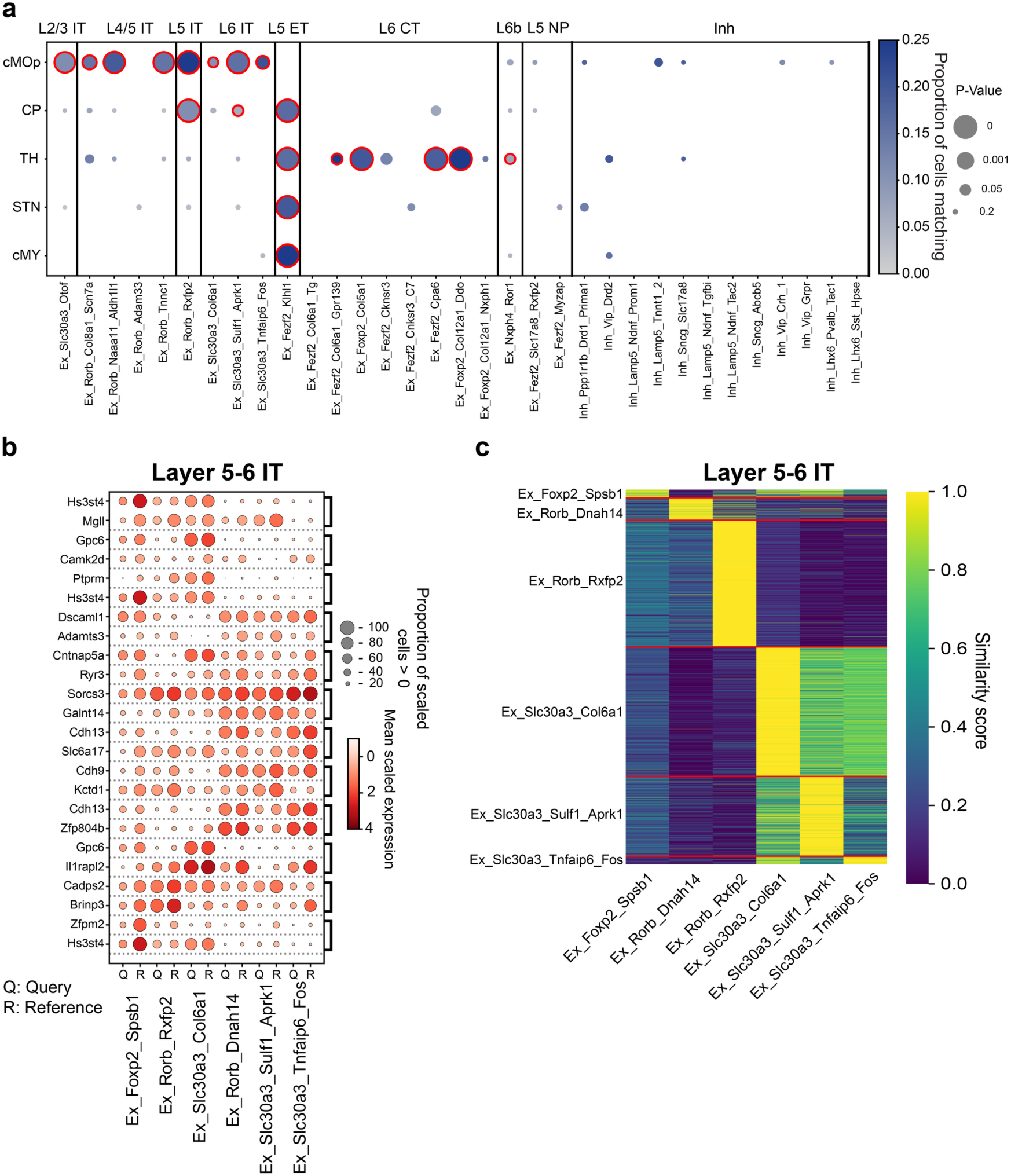
Fine-grained cell type predictions of neuronal populations in AC presynaptic Synapse-seq experiment. **a**, Dotplot of projection matching for each cluster type expressing at least 200 VTs (full statistical results reported in Supplementary Table 10). **b**-**c**, Dotplot (**b**) and heatmap (**c**) of scaled correlations confirming the identity of layer 5-6 IT cluster types. Markers for both were computed on a sampled reference, matching the composition of the predicted query dataset.

**Extended Data Fig. 9:**
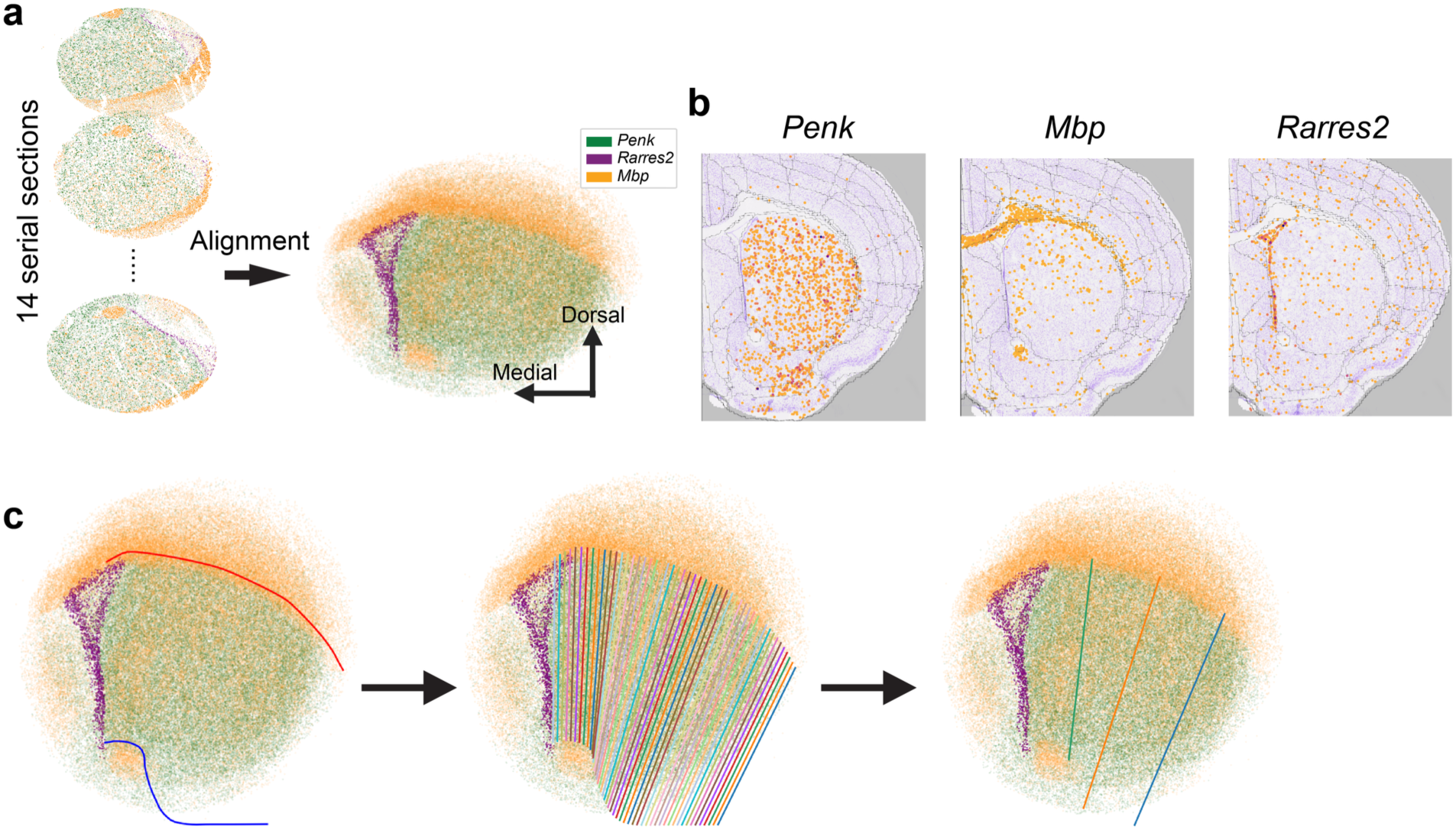
Alignment of serial striatal sections to quantify corticostriatal projections from AC. **a**, Alignment of 14 serial Slide-seq arrays (left) showing resulting regional marker expression (right). **b,** Expression of alignment markers mapped in the whole-brain atlas^1^. **c**, Definition of striatal subregions: drawing bounding curves (left), partitioning based on depth (middle), and selecting zones for analysis (right).

**Extended Data Fig. 10:**
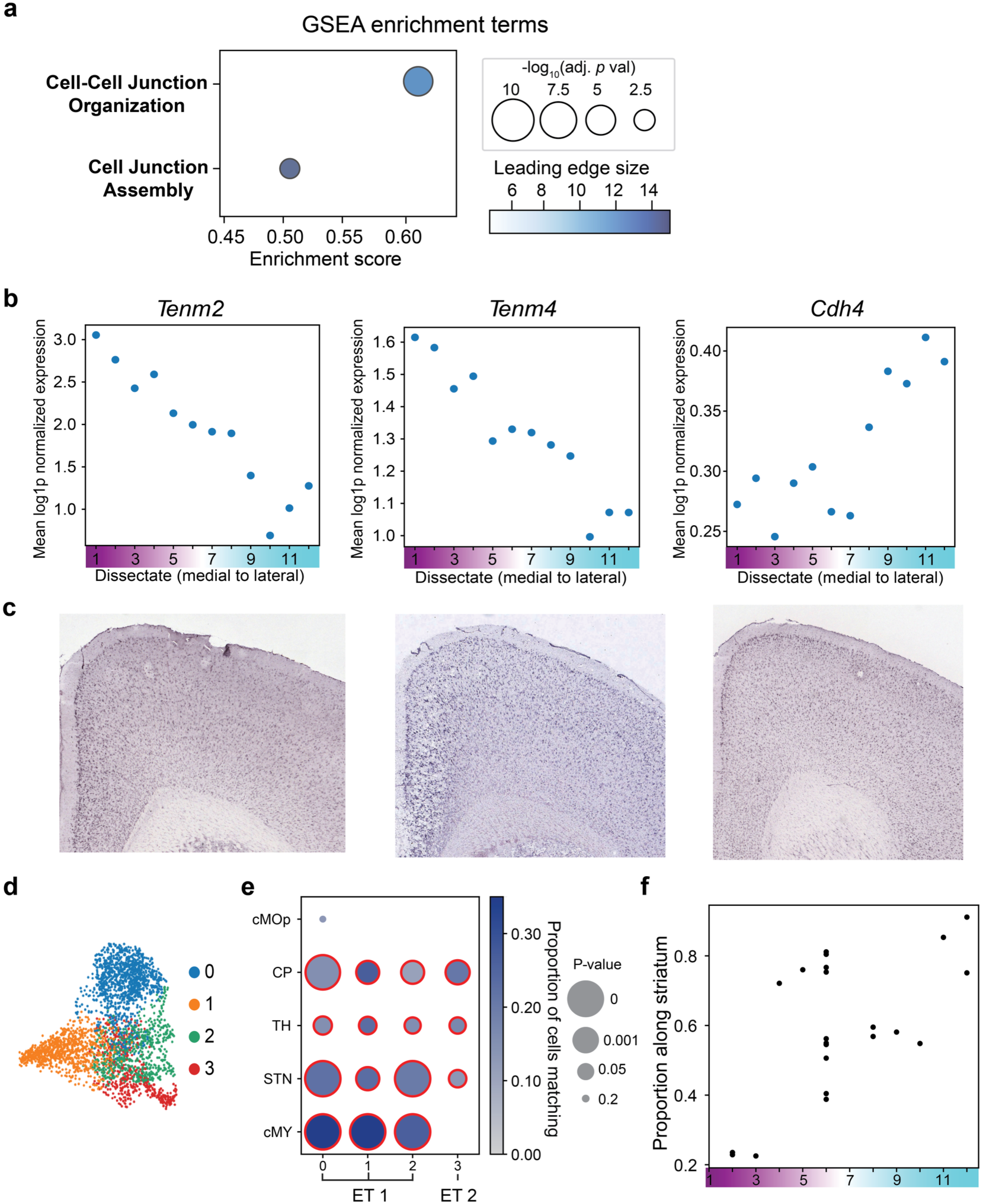
Extended analyses of L5 ET corticostriatal projections. **a**, Dotplot showing significant GSEA results identifying gene sets enriched at one end of the cortical dissection axis shown in Fig. 4a. **b**,**c**, Expression of three genes correlating with the medial-lateral axis (**b**) and corresponding in situ hybridization (Allen Brain Atlas) (**c**). **d**, UMAP embeddings of layer 5 ET cells, colored by de-novo cluster assignment. **e**, Projection matching of L5 ET subclusters to targets; groupings into ET 1 and ET 2 indicated on x-axis. **f**, Correlation between cortical medial-lateral position (Fig. 4a) and striatal projection axis (**Extended Data** Fig. 9c) for ET 1 neurons (*n* = 23 VTs, projection score > 0.9).

**Extended Data Fig. 11:**
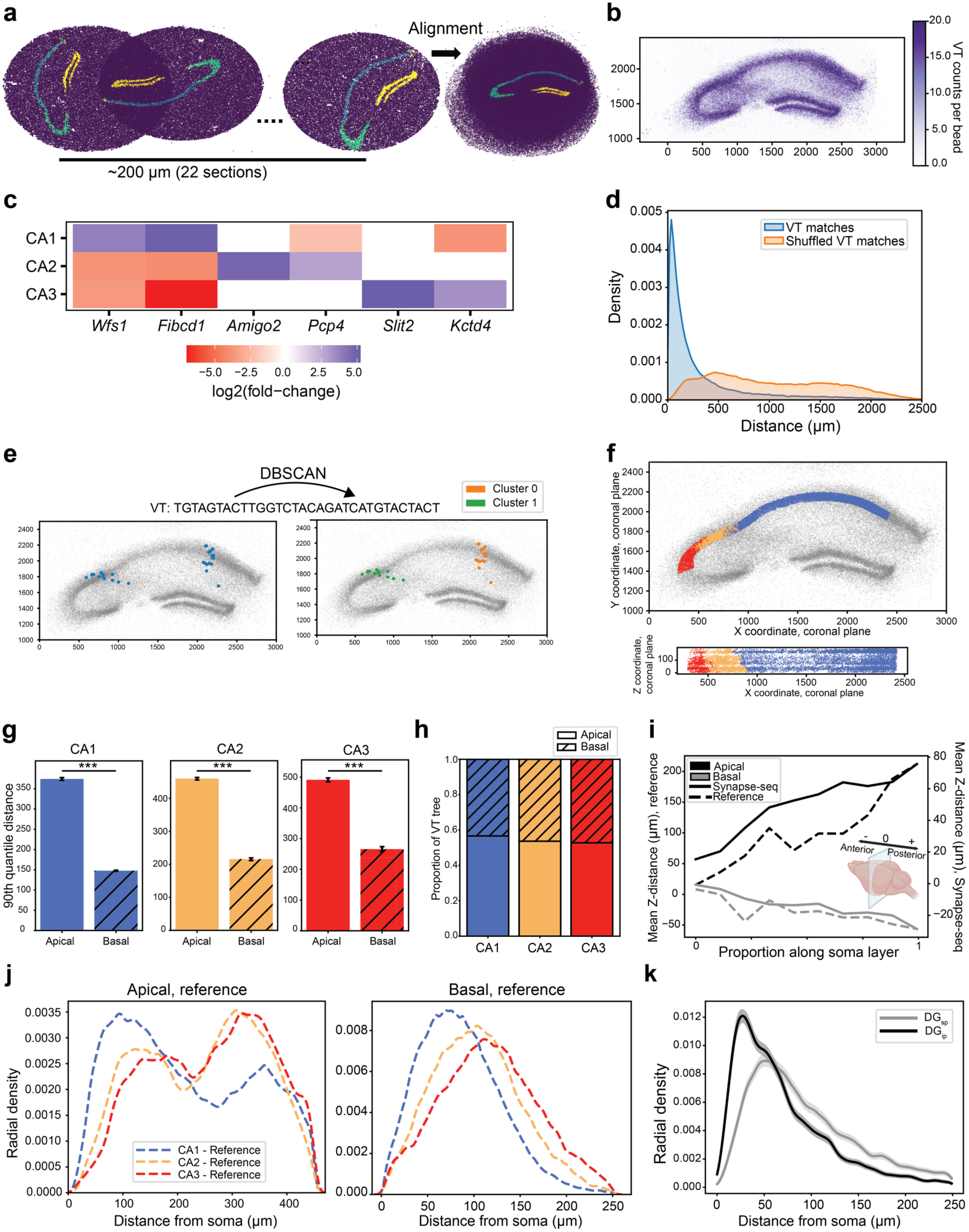
Processing and benchmarking of postsynaptic barcoding data for dendritic morphological analysis. **a**, Alignment of 22 serial Slide-seq sections (∼200 μm in anterior-posterior z distance) into a single plane using pyramidal (blue/green) and dentate granule (yellow) de novo cluster landmarks. **b**, Per bead VT UMI counts in collapsed (x,y) coronal space (axis distances in μm). **c**, Marker expression heatmap across subfields. **d**, Pairwise bead distance distributions for observed (blue) vs. shuffled (orange) data. **e**, Correction of VT “collisions” via DBSCAN (axis distances in μm). **f**, RCTD-annotated estimated soma locations in both x-y space (top, coronal plane) and x-z space (bottom) (axis distances in μm). **g**, Barplots depicting the 90th quantile of distance between apical or basal VT matches and their cell bodies, separated by hippocampal neuronal type (mean ± 95% confidence interval; *** p < 0.001, bootstrap test). **h**, Barplots depicting the proportion of VT trees found in the apical compared to the basal layers across subfields. **i**, Dendritic tree orientation in anterior-posterior z axis vs. soma position compared to reference^46^. **j**, Radial density analysis of apical and basal dendrites in the reference dataset^46^. **k**, Radial density analysis of DG_sp_ and DG_ip_ with 95% confidence intervals shaded in gray.

**Extended Data Fig. 12:**
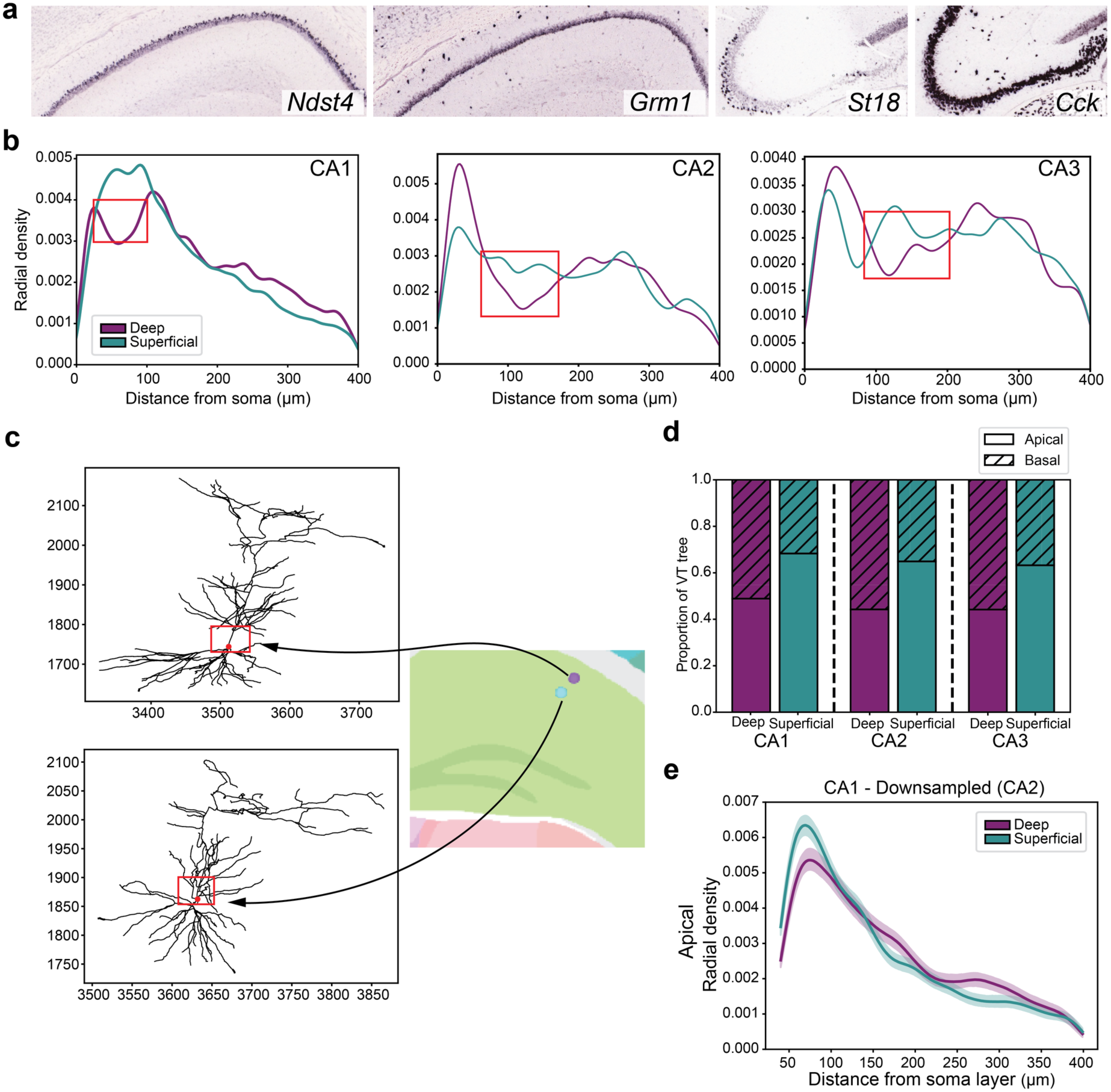
Transcriptomic and anatomical evidence of superficial-deep heterogeneity across hippocampal subfields. **a**, Expression of canonical superficial and deep markers in CA1 (*Ndst4*, *Grm1*) and CA3 (*St18*, *Cck*) from Allen Mouse Brain Atlas *in situ* hybridization data^52^. **b**, Radial density analysis of apical VT matches for superficial and deep somas, separated by hippocampal subfield. Red boxes indicate the presence (deep) or absence (superficial) of a “neck” of reduced branching in the proximal apical dendrite. **c**, Dendritic reconstructions^46^ of representative deep (top, neuron ID: 202410_013) and superficial (bottom, neuron ID: 202829_095) cells located in the CA1, with their soma locations (right). **d**, Barplots depicting the proportion of VT trees found in the apical compared to the basal layers between both superficial and deep cells, across subfields. **e**, Radial density analysis of the apical CA1 layer, downsampled to have the same number of total VT matches in the apical layer as CA2. 95% confidence intervals shaded.

## Methods

### Experimental Methods

#### Mice

All *in vivo* experiments were performed in male C57BL/6J at ages P90-P120, purchased directly from The Jackson Labs (Strain #: 000664). All mice were housed on a 12-h light/dark cycle between 68 °F and 79 °F and 30–70% humidity. All animal work was approved by the Broad’s Institutional Animal Care and Use Committee (IACUC).

#### Primary cortical neuron culture

Rat primary cortical neurons (Thermo Fisher Scientific) were thawed and plated at a density of 40,000–60,000 cells per well on Poly-D-Lysine-coated coverslips in plating medium (NBactiv4 supplemented with N-2, 5% heat-inactivated FBS, and gentamicin). On day 5, cultures were transitioned to serum-free synapse maturation medium (NBactiv4 with N-2 and gentamicin). Transfections were performed at DIV9–14 using the CalPhos Mammalian Transfection Kit (Takara). Neurons were incubated for 45 minutes with a DNA-calcium phosphate precipitate formed from 1.6 µg total DNA (including pUC19 filler) per well. Following a brief wash in HEPES-NaCl buffer, neurons were returned to their original conditioned media and analyzed 24 hours later.

#### *in vitro* and *in situ* detection of Synapse-seq components

Primary neurons were fixed in 4% paraformaldehyde, dehydrated in an ethanol (EtOH) gradient, and treated with protease K digestion for signal enhancement. Mice injected with the postsynaptic targeting system were perfused in 10mL PBS then 10mL 4% PFA, brains were extracted, cryosectioned into 20 μm-thick sections, mounted on Superfrost Plus slides, and dehydrated in an EtOH gradient. Hybridization Chain Reaction (HCR RNA-FISH v3.0, Molecular Instruments) was used with a custom *mScarlet*-B4 probe according to manufacturer’s instructions and modified as published^64^. Mice injected with the presynaptic targeting system were perfused in HBSS and brains were snap-frozen in OCT, cryosectioned into 12 μm-thick sections, mounted on Superfrost Plus slides, fixed in 4% paraformaldehyde, dehydrated in an EtOH gradient, and treated with hydrogen peroxide. RNAscope Fluorescent Multiplex Kit v2 (ACDBio) with probe mScarlet-C1 (Cat No. 572421) and TSA Plus Cy5 dye (Akoya Biosciences) was used according to manufacturer’s instructions, besides removing the Protease IV treatment step to improve preservation of targeting protein signal. For dual smFISH-IHC experiments, samples were then washed in PBS for 10 min, blocked in 4% NGS in PBS-T (0.3% Triton-X) for 1 hour at RT, treated overnight with either anti-GFP polyclonal antibody (1:1000, Abcam ab13970) (Fig. 1c-g, Extended Data Fig. 1n) or anti-Bassoon monoclonal antibody (1:400, Enzo SAP7F407) (Fig. 1h,i), washed 3 x 5 min in PBS at RT, then incubated with secondary antibodies in 4% NGS in PBS-T for 2 hours at RT with gentle agitation (1:1000, goat anti-Chicken IgY (H+L) Alexa Fluor 488, invitrogen A-11039; 1:400, goat anti-mouse (H+L) Alexa Fluor 405, invitrogen A-31553). Samples were co-stained with DAPI or Hoechst (except with 405 secondary) and mounted in Fluoromount-G (invitrogen) or ProLong Gold Antifade Mountant (Thermo Fisher).

#### Microscopy

##### Acquisition

Primary neuron cultures, the RSC (Extended Data Fig. 1g-h), and the SSp (Extended Data Fig. 1j-k) were imaged on an Andor CSU-X spinning disk confocal system coupled to a Nikon Eclipse Ti microscope equipped with an Andor iKon-M camera (20x or 60x oil objectives). All other imaging was performed on the Zeiss LSM 900 confocal microscope (20x or 63x oil objectives).

##### Quantification

RNAscope puncta density and colocalization with targeting protein or Bassoon were quantified using ImageJ v1.53t (Fig. 1, Extended Data Fig. 1). Automatic thresholding (calculated for a stem-loop-positive sample from each experimental batch, then applied to stem-loop-negative controls within the same batch using the algorithms listed below), binary masks, dilation, watershed, and analyze particles (size restriction imposed to detect synaptic puncta while excluding larger objects, such as retrograde or transsynaptic labeling of cell bodies^65^) were used to calculate puncta density, followed by imageCalculator() to calculate percent of puncta with colocalized protein. The following thresholding algorithms were used: LGd: Moments for GFP, Default for mRNA; RSC: Otsu for GFP, Default for mRNA; VPM: Otsu for GFP, Li for mRNA; CP: Default for GFP, Otsu for mRNA; HPC: Default for GFP, Moments for mRNA. Unpaired two-tailed Welch’s t-tests were performed as indicated with Welch-Satterthwaite degrees of freedom reported (Fig. 1e,g,m). Profile analysis (Fig. 1k) in Zeiss ZEN blue v3.7.5 software was performed to compute the fluorescence intensity (arbitrary units, a.u.) of mRNA, targeting protein, and DAPI along a perpendicular line through the soma layer, spanning the full apical and basal fields of the CA1 as indicated by the boundary of GFP targeting protein signal. A 60 μm soma layer was manually segmented out using the fluorescence intensity of the *mScarlet* mRNA and DAPI signal.

#### Cloning and packaging of barcoded AAV

##### Generation of barcoded plasmid

The barcoded plasmid was generated by digesting the backbone *pAAV-CAG-MVE-xrRNA-mScarlet-24xPP7* with XhoI and SphI-HF (New England Biolabs). The insert was amplified using Q5 High-Fidelity DNA Polymerase (New England Biolabs) with the *WPRE_ssDNA_32bp_10Xbc* forward primer and *Sph1_reverse_primer_short* (Supplementary Table 8). Cycling conditions were: 98°C for 30 s, followed by 24 cycles of 98°C (10 s), 65°C (20 s), and 72°C (60 s), with a final extension at 72°C for 5 min. Following DpnI digestion and SPRIselect purification (Beckman Coulter), the vector and insert were assembled (1:3 ratio) using HiFi DNA Assembly Master Mix (New England Biolabs) at 50°C for 1 hour. The assembly reaction was cleaned up by sequential incubation with QuickCIP and T5 Exonuclease (New England Biolabs), followed by a final 1.8x SPRIselect purification.

##### AAV production and titering

HEK293T/17 cells (ATCC) were transfected with barcoded plasmid, adenovirus pHelper, and AAV2/9 (rep-cap) at a 1:2:4 ratio (39.93 µg total). Media and cell lysates were harvested approximately 60 hours post-transfection and purified as described^66^. To determine titers, purified AAVs were treated with Turbonuclease (37°C, 1 h) to remove unencapsidated DNA, followed by Proteinase K digestion (56°C, 2–16 h) to release viral genomes. Viral genomes were quantified by droplet digital PCR (ddPCR) using a QX100 system (Bio-Rad) with primers and a probe targeting the ITR (Supplementary Table 8), using an annealing/extension temperature of 58°C. pAAV-CAG-MVE-xrRNA-mScarlet-24xPP7 (unbarcoded), pAAV-CAG-MVE-xrRNA-mScarlet (stem-loop-negative control), and pAAV-hSyn1-SYP-sfGFP-tdPCP vectors were packaged by Vector Biolabs (PA, USA). pAAV-hSyn1-PSD95.FingR-sfGFP-ZF-tdPCP and pAAV-hSyn1-GPHN.FingR-sfGFP-ZF-tdPCP were packaged by Boston Children’s Hospital Viral Core (MA, USA). Titers ranged from 6.4e12 - 6.40e13 viral genomes per mL.

#### Sequencing the barcoded AAV pool

##### Viral vector digestion

To quantify barcode diversity, 2 µL of viral vector was incubated with 100 µL of fresh DNAse I solution (50 U/mL DNAse I in 2 mM CaCl₂, 10 mM Tris-HCl pH 7.6, 10 mM MgCl₂) at 37°C for 1 hour to degrade non-encapsidated DNA. The reaction was quenched with 5 µL of 0.5 M EDTA (invitrogen) and heat-inactivated at 70°C for 10 minutes. Capsids were then digested by adding 120 µL of Proteinase K solution (100 µg/mL Proteinase K in 1 M NaCl, 1% N-lauroylsarcosine) and incubating at 50°C for 2 hours, followed by inactivation at 95°C for 10 minutes.

##### UMI tagging and library preparation

Digested viral genomes were tagged with a unique molecular identifier (UMI) via a single-cycle PCR. The reaction contained 24 µL of digested vector, 1 µL of 10 µM primer OL1431 (Supplementary Table 8), and Kapa HiFi HotStart ReadyMix (Roche). Cycling conditions were: 98°C for 2 min, followed by a single cycle of 98°C (20 s), 62°C (30 s), and 72°C (20 s), with a final extension at 72°C for 5 min. The resulting UMI-tagged product was then amplified for sequencing using primers OL869 and OL10 (Supplementary Table 8). This second PCR used an initial denaturation at 98°C (2 min), followed by 8 cycles of 98°C (20 s), 62°C (15 s), and 72°C (15 s), and a final extension at 72°C (5 min).

#### Stereotaxic surgery for AAV delivery

AAV vectors for each Synapse-seq component were mixed 1:1 by volume and stereotactically injected into the target regions listed below. All injections were delivered at a rate of 1 nL/s, and the capillary was left in place for 5-10 minutes post-injection to prevent backflow. Mice were perfused after 1-5 weeks of viral incubation.

Table of AAV injections and their volumes per brains region:

| Region | Coordinates (relative to bregma and skull) | Volume of virus mix |
| --- | --- | --- |
| Midbrain<br>(substantia nigra, SN) | -2.92 mm anterior-posterior, -1.15 mm<br>mediolateral, -4.64 mm dorsoventral | 100 nL |
| Primary Visual<br>Cortex (VISp) | -2.8 mm anterior-posterior, +/- 2.6 mm<br>mediolateral, -1 mm dorsoventral | 50 nL |
| Ventral<br>posteromedial<br>nucleus (VPM) | -1.7 mm anterior-posterior, + 1.75 mm<br>mediolateral, -3.25 mm dorsoventral | 150 nL |
| Hippocampus<br>(Fig. 1j-l) | -1.6 mm anterior-posterior, +/- 1.75 mm<br>mediolateral and -1.25 mm dorsoventral | 150 nL |
| Anterior Cortex<br>(AC) | +1.8 mm anterior-posterior, + 1 mm mediolateral<br>and 1.3 dorsoventral (medial); +1.8 mm anterior-posterior, + 2 mm mediolateral and 1.3<br>dorsoventral (lateral) | 100 nL each |
| Hippocampus<br>(Figs. 5-6) | -2.2 mm anterior-posterior, +/- 1.3 mm<br>mediolateral, -2.0 mm dorsoventral | 150 nL |

#### Single-nucleus RNA sequencing

Mice were perfused with HBSS (Thermo Fisher Scientific) and extracted brains were snap-frozen in OCT (Tissue-Tek). To localize the injection site, 14 µm cryosections were fixed in 70% ethanol, stained with DAPI, and examined for GFP fluorescence. A 1 mm biopsy punch was collected from the identified region, cryosectioned into a 300–500 µm slice, and the GFP-positive injection site was microdissected into RNase-free tubes. Samples were stored at −80°C for up to 24 hours.

Nucleus isolation was performed as previously described^67^, counted via flow cytometry, and loaded into the 10x Chromium system using the Single Cell 3′ v3.1 kit. Gene expression libraries were constructed according to the manufacturer’s protocol. Targeted viral transcript sequencing libraries were prepared in parallel as described below.

#### Bulk RNA extraction

Tissue samples were obtained by cryostat microdissection at −20°C, and total RNA was extracted using the RNeasy Mini Kit (Qiagen). RNA integrity was assessed using an Agilent Bioanalyzer with the RNA 6000 Nano Kit. Reverse transcription was performed using Maxima H Minus Reverse Transcriptase (Thermo Fisher Scientific) with primer SL59 (Supplementary Table 8), which contains a 10-nucleotide unique molecular identifier (UMI) and a sequence complementary to the Synapse-seq barcode construct. The resulting RNA/cDNA duplexes were purified using AMPure XP beads (Beckman Coulter).

To enhance target specificity, a first round of nested PCR was performed using >2 ng of purified cDNA, Kapa HiFi HotStart ReadyMix (Roche), and primers OL10 and SL61 (Supplementary Table 8). Cycling conditions were: 98°C for 2 min, followed by 13 cycles of 98°C (20 s), 67°C (15 s), and 72°C (15 s), with a final extension at 72°C for 5 min. The resulting product was purified using SPRIselect via a double-sided (0.6x/0.4x) then left-sided (1.2x) size selection, and used as input for the final targeted viral transcript amplification as below.

#### Slide-seq assays

Fresh frozen brains were trimmed to the *mScarlet*-expressing injection site, and 5-20 µm consecutive coronal sections were transferred onto Slide-seq arrays. Slide-seqV2 was performed as described^32^ with modifications: RNA hybridization was conducted in 200 µL hybridization buffer (6x SSC, 10% sarkosyl, 2 U/µL Lucigen NxGen RNAse inhibitor) for 20 minutes at room temperature. cDNA libraries were amplified using Terra Direct PCR Mix (Takara) with the Truseq PCR handle primer, SMART PCR primer, and OL1364 primer (Supplementary Table 8). Targeted viral transcript sequencing libraries were prepared in parallel as described below.

#### Targeted viral transcript amplification

Viral transcripts were enriched from >2 ng of the above barcoded cDNA libraries (snRNA-seq, Slide-seq) or nested PCR product using Kapa HiFi HotStart ReadyMix and primers OL10 and OL869 (Supplementary Table 8). Amplification was performed with an initial denaturation at 98°C (2 min), followed by 13–18 cycles of 98°C (20 s), 67°C (15 s), and 72°C (15 s), and a final extension at 72°C (5 min). PCR products were purified using SPRIselect beads (Beckman Coulter) via a double-sided size selection (0.6x/1.2x) and assessed on an Agilent Bioanalyzer High Sensitivity DNA chip.

#### Sequencing

Samples were pooled and sequenced on the Illumina NextSeq or NovaSeq flowcells with a read structure of 42 bases Read 1, 10 bases i7 index read, 10 bases i5 index read, 80 bases Read 2. Each dial-out sample received approximately 50-150 million reads. Slide-seq gene expression libraries received 200-400 million reads, corresponding to 3,000-5,000 reads per bead. Single-nucleus RNA-sequencing gene expression libraries received 60,000 reads per cell.

### Computational Methods

#### Synapse-seq Data Preprocessing

##### Viral transcript detection and processing

Reads containing UMI-barcoded viral transcripts (VTs) were filtered by fuzzy matching (Hamming distance ≤ 3) to the flanking constant sequences and VT structure. To generate a high-confidence barcode whitelist, two independent viral aliquots were sequenced; singleton VTs were collapsed to more abundant sequences (Hamming distance 1), followed by weighted graph-based deduplication using UMIcollapse v1.0.0 (k=2). This process identified and removed 6.6% of observed VTs—primarily low-abundance sequences supported by single UMIs—resulting in a final validated VT whitelist.

##### Single-nucleus RNAseq processing

Raw BCL files were demultiplexed and aligned to a GRCm39 reference augmented with the mScarlet sequence using Cell Ranger v7.2.0. Unfiltered expression matrices were denoised with CellBender v0.3.0 (FPR = 0.01, learning rate = 5e-4) to remove ambient background; only droplets containing at least one mScarlet transcript were retained. Data were processed in Scanpy v1.9.3 (2,000 highly variable genes, 80 PCs, Leiden resolution = 3). Nuclei were flagged as low-quality or doublets if they were identified by Scrublet v0.2.3 (expected rates: 0.15 VISp, 0.20 AC), resided in clusters with ≥25% doublet calls or mixed canonical markers (Supplementary Table 9), or if ≤70% of their 10-nearest neighbors belonged to the same subclass. This filtering removed 33,889 (34%) and 60,491 (27.6%) nuclei from the VISp and AC datasets, respectively.

##### Slide-seq spatial transcriptomics data processing

Slide-seq gene expression libraries were demultiplexed, aligned to the augmented GRCm39 reference, and spatially placed using SlideSeqTools with default settings^31^ (https://github.com/MacoskoLab/slideseq-tools).

##### Processing of viral transcript counts

We developed a unified pipeline to map targeted reads to the validated VT whitelist. For bulk and snRNA-seq data, reads were filtered by fuzzy matching both pre- and post-barcode constant sequences (Hamming distance ≤ 3). VTs were assigned to the whitelist if they fell within a Hamming distance of 1; ambiguous matches equidistant to multiple whitelist sequences were discarded. For snRNA-seq, cell barcodes were corrected to the 10x Genomics whitelist (Hamming distance ≤ 1). For Slide-seq, beads were merged based on sequence similarity and spatial proximity (10 µm), and VTs were identified by matching the post-constant sequence (Hamming distance ≤ 5) and the whitelist (Hamming distance ≤ 1).

Across all modalities, UMI-VT combinations were graph-deduplicated using UMICollapse^68^ (k=1). For UMIs associated with multiple VTs, the identity was assigned based on maximum read support. To remove PCR recombination artifacts, molecules were filtered based on read depth: bulk data utilized a 90th quantile reads/UMI cutoff, while snRNA-seq and Slide-seq libraries used an inflection point determined by elbow plot analysis of reads per UMI versus total observed molecules.

#### Cell type assignment of snRNA-seq data in VISp and AC experiments

##### Label transfer to a whole mouse brain atlas

Cell types were assigned by transferring labels from a whole-mouse-brain single-nucleus RNA-seq atlas^1^ to the VISp and AC snRNA-seq datasets. Similar to previously described approaches^2^, we utilized a hierarchical classification strategy (class, subclass, cluster) based on pairwise marker genes and k-nearest neighbor (KNN) classifiers using cosine distance. This approach compared query cells to a sub-sampled reference dataset rather than centroids, and by performing a secondary KNN classification among the top 15 predicted clusters to refine assignments. To validate predictions, the reference atlas was subsetted to clusters present in the query (>30 cells), and expression data were log1p-normalized. Marker genes were identified using COSG (class level)^69^ or Wilcoxon rank-sum tests (subclass/cluster levels) and visualized via dotplots with independent scaling to account for sequencing depth differences. Label transfer efficacy was further quantified by computing a similarity score, defined as the mean cell-wise correlation between a query cell and reference subclass cells using the top 20 positive and negative subclass markers. Code used for the label transfer procedure can be found at: https://github.com/MacoskoLab/pknn_repo/tree/main.

##### Quantification of transcriptomic impact of AAV transduction in VISp dataset

To benchmark viral perturbation, we mapped Synapse-seq and Sindbis-transduced^24^ profiles to a whole-brain reference atlas^2^ using the MapMyCells label transfer approach^70^ optimized for the reference. Raw Sindbis reads (SRR9858304, SRR9858308) were re-aligned (Cell Ranger v7.2.0, GRCm39) and denoised with CellBender v0.3.0 (FPR=0.01, learning-rate=5e-4). Analyses were restricted to cells present in the original dataset^71^ with > 200 umi and < 10% mitochondrial gene UMIs. Transduction was defined as >= 10^2.8^ viral barcodes for Sindbis and concurrent detection (>= 1 count) of viral mRNA and dialout barcodes for AAV (the same criterion for projection analysis). Cell-class marker genes were identified using COSG^69^; expression profiles were log1p-normalized and scaled independently in Scanpy for visualization.

#### Computational matching of presynaptic barcodes

To distinguish true projections from technical noise, we calculated a weighted “VT score” for each match that accounts for viral barcode collisions and ambient RNA contamination. First, we modeled the probability of barcode collisions—where a single viral sequence transduces multiple cells—using a binomial distribution: 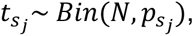 Here, 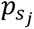 is the prevalence of sequence *s_j_* in the viral pool, and *N* (total cellular transductions) was estimated by dividing the count of observed injection-site VTs by the detection sensitivity (the proportion of projection-site VTs detected at the injection site). Matches were weighted by the probability that a barcode was attributable to the observed cell, given it was transduced, rather than a collision event 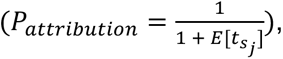 effectively down-weighting abundant barcodes. Second, we accounted for ambient contamination by incorporating the probability that a VT originated from the background *P_ambient_*. This was derived from a likelihood ratio of UMI counts (*U_i_*_,*j*_) in mScarlet-positive neurons versus empty droplets, scaled by cell-specific background fraction prior (*b_i_*)estimated by CellBender. The final VT score was computed as the product of the attribution probability and the transduction probability (1 – *P_ambient_*).

To assess the robustness of these scores, we performed jackknife resampling, iteratively holding out 10% of the data to quantify their variability. Statistical significance for cell-type-to-region projections was determined using a one-sided non-parametric test against a “glial null” distribution, generated by resampling glial cells to match the sample size of the query cell type. The final significance value was reported as the mean p-value across jackknife iterations. For projection site analysis, we subsetted excitatory neurons to cortical subtypes, excluding the L2/3 IT subtype in the VISp experiment due to low transduction rate and low number. Full projection results and p-values can be found in Supplementary Table 10.

##### Collision rate estimation

To model the rates of collision for different numbers of total cellular transductions, *N_c_*, we ran simulations for a range of hypothetical *N_c_* (1,000-1,000,000 cells). For each *N_c_*, we sampled VTs using the same binomial described above, assuming an average of 6.5 VTs per cell. From these distributions we computed the expected proportion of total molecular collisions, along with the variance in collision probability across representative VTs (Extended Data Fig. 4c).

##### Relative noise contribution

To quantify false-positive matches arising from ambient RNA, we combined cell-specific background fractions (*Bg_RNA_* = 0.077) estimated by CellBender with the projection matching rate observed in empty droplets (*r_empty_* = 0.0435). The expected number of ambient matches within the glial population (*n_glia_* = 75,517) was calculated as *n_glia_* ∗ *Bg_RNA_* ∗ *r_empty_* = 254.32. Comparing this value to the total observed glial matches (2,909) indicates that ambient contamination accounts for approximately 9% of false-positive calls.

#### Presynaptic spatial data processing and analysis

##### Slide-seq array alignment and VT localization

Serially sectioned arrays from the thalamus (15 sections), striatum (14 sections), and superior colliculus (16 sections) were aligned using regionally specific molecular markers and viral barcodes. Alignment was performed using an open-source pipeline (https://github.com/mukundraj/broad-registration/tree/main) refined by manual linear shifts and rotations (Extended Data Figs. 6, 9). To analyze the spatial distribution of projections, VTs originating from the same cell were computationally collapsed into single data points. Matches were attributed to a single source cell if all spatially localized VTs of a given sequence fell within a dynamic radius (146 μm * number of capture locations, max 730 μm) of their combined centroid; matches failing this criterion were discarded as likely barcode collisions or recombination artifacts. Thresholds were chosen based on distribution of pairwise distances of beads with the same VT. This filtering retained 90.7% and 91.1% of VTs in the striatum and thalamus, respectively.

##### Visual cortex corticothalamic analysis

To subset viral transcripts to specific nuclei, the dorsal lateral geniculate nucleus (LGd) and lateral posterior nucleus (LP) were manually outlined based on marker gene expression (Fig. 2f). The LGd was defined by the ventral LGv (*Gad2*), the inferior white matter tract (low UMI, *Pvalb*-negative), and a medial boundary marked by higher-order nuclei (*Necab1*) and the dentate gyrus (*Prox1*). The LP was delineated by *Necab1* expression, bounded inferiorly by the posterior complex (*Tnnt1*-positive, *Necab1*-negative) and superiorly by the *Gad2*-positive dentate gyrus.

Laminar depth of Layer 6 corticothalamic (CT) subtypes was quantified using spatial coordinates from a whole-brain reference atlas^1^ mapped to the Common Coordinate Framework. To normalize depth, we fit a characteristic superficial boundary curve (third-degree polynomial) to Layer 4/5 IT neurons (*Ex_Rorb_Naaa11_Aldh1l1*) in the VISp or SSp. The minimum distance from each mapped L6 CT bead (mapping score ≥ 0.15) to this curve was calculated, with 95% confidence intervals estimated via 10,000-iteration bootstrap resampling weighted by mapping confidence.

##### Anterior cortex corticostriatal analysis

Cortical regions were computationally excluded from striatal arrays by demarcating boundaries at the corpus callosum (*Mbp*) and medial ventricle (*Rarres2*). We defined a coordinate space based on bounding curves drawn at the corpus callosum and anterior commissure (Extended Data Fig. 9). Bead coordinates were transformed into distances along (*r*), and between (*i*), these curves, and the striatum was subdivided into equidistant sections based on *r*. To visualize projection topographies, we fit a Gaussian kernel density estimate on a meshgrid of all striatal matching VTs for a given cell type (weighted by projection score) and plotted the resulting contours over the aligned arrays.

To quantify topographic projections to the medulla, cortical dissections were grouped into medial, mid, and lateral bins (Fig. 4a). Medullary dissections were assigned pseudocoordinates, and the mean y-coordinate (dorsal-ventral axis) was calculated for each cortical group based on its medullary matches, weighted by the attribution probability (*P_attribution_*) for any given VT. VTs were filtered if supported by a single UMI or if <80% of UMIs fell into dissections sharing a y-coordinate (>80% of VTs met these criteria). We computed 95% confidence intervals on mean medullary matching, using bootstrap re-sampling with replacement, for each cortical grouping. We computed the bootstrapped t-test difference in means (scipy.stats.ttest_ind) between each cortical region, using 10000 resamplings.

To identify genes correlated with the medial-lateral cortical axis (Fig. 4e), snRNA-seq libraries were grouped by dissection and mean log1p-normalized expression was calculated. Genes expressed in fewer than 500 cells were filtered. Pearson correlations were computed between mean expression and the integer-encoded dissection position (1 to 12). Genes with correlation magnitude at least 0.65 were scaled across dissectates and plotted. GSEA was performed using the gseapy v1.1.3 prerank module on correlation-ranked features.

To relate medullary targeting to striatal collateralization, we calculated a striatal enrichment score. VTs were categorized by their medullary target location (ventral: y<1; mid: y=1; dorsal: y>1). Striatal match scores for these VTs were directionalized (+1 for ventral, 0 for mid, -1 for dorsal medullary targets), and the mean directionalized score was computed for each striatal region. We performed a permutation test on the difference-of-means t-test (scipy.stats.ttest_ind) between the scaled VT distributions of the two most medial versus two most lateral striatal sections.

#### Postsynaptic data processing and analysis

##### Alignment and cell type deconvolution

De novo clustering of 22 serial Slide-seq arrays was performed using Seurat v5 (dims=1:10, res=1.5). Clusters expressing pyramidal (*Neurod6*) or dentate granule (*Prox1*) markers served as landmarks to align arrays into a common (x,y,z) coordinate space via an open-source registration pipeline (https://github.com/mukundraj/broad-registration/tree/main). Beads were assigned cell types using Robust Cell Type Decomposition (RCTD, spacexr v2.2.1)^72^ in ‘doublet’ mode against a reference atlas^1^. The reference was subsetted to clusters with >500 nuclei in the dorsal hippocampal subfields: CA1 (*Ex_Crym_Chrna5*), CA2 (*Ex_Crym_Nuak2*), CA3 (*Ex_Crym_Csf2rb2*, *Ex_Crym_Cldn22*, *Ex_Crym_Plagl1_1*, *Ex_Crym_Plagl1_2*), and DG (*Ex_Crlf1_Ntn5_1*, *Ex_Dsp_Moxd1_1*, *Ex_Dsp_Moxd1_2*, *Ex_Dsp_Moxd1_3*, *Ex_Dsp_Gsg1l*). Beads were filtered for <5% mitochondrial reads. Subfield-specific markers were identified using the limma implementation of the Wilcoxon Rank Sum test (logfc.threshold=0, min.pct=0.1) on RCTD singlets.

##### Dendritic reconstruction by VT trees

Hippocampal soma layer boundaries and midlines were segmented using the cgrid package (https://github.com/MacoskoLab/Macosko-Pipelines/blob/main/tools/cgrid/cgrid.R). To resolve VT collisions, spatially distinct barcode occurrences were clustered using DBSCAN (eps=300 μm); only VTs detected on ≥4 beads were retained. For each resulting “VT tree,” the soma position was estimated by fitting a kernel density estimator (bandwidth=100) to VT coordinates on a 20×20×20 voxel grid and optimizing for the maximum density within the soma layer. Trees were retained if they contained ≥4 UMIs in the soma layer and ≥5 UMIs in synaptic layers. Cell type was assigned to each VT tree by majority vote of the 10 beads nearest the soma. Total cell counts were estimated using a correction factor of ∼1.9 VT trees per cell, derived from Slide-tags performed on two of the 22 serial sections, where nuclear profiles were intersected with proximal (100 µm) VT trees.

##### Morphological annotation and benchmarking

VT matches were mapped to a polar coordinate system (*r*, *ϕ*) centered on the estimated soma, where *r* is the three-dimensional Euclidean distance from the soma (filtered to *r*<500 μm) and *ϕ* is defined relative to the local perpendicular of the soma layer midline. To adjust for angular differences observed along the soma layers (e.g. CA3 trees were less perpendicular to the soma layer than CA1 trees), VT trees were rotated as needed to ensure proper polar coordinate alignment. Each VT match was classified as above or below the soma based on its signed projection onto the VT tree’s top principal component (PCA); apical versus basal was then assigned under the assumption that apical dendrites are longer than basal dendrites (95% quantile of distance from soma). Assignments were manually verified. Reference reconstructions of hippocampal neurons (https://mouse.digital-brain.cn/projectome/hipp)^46^ were processed for benchmarking by filtering for CA1, CA2, and CA3 pyramidal cells that were located near our coronal cross-section (7000-7500 μm in the anterior-posterior z-axis). After manually selecting for pyramidal morphologies, we retained full dendritic reconstructions of 111 neurons of interest (94 CA1, 9 CA2, 8 CA3). To emulate Synapse-seq measurements, we converted the graph encoding of .swc files into an undirected set of points randomly sampled from each 1 μm segment along the dendritic branch. Manual drawing of polygons delineating the apical and basal layers of these reconstructions were performed in order to annotate points. We supplemented these data with wild type dentate granule cells^73^ from https://neuromorpho.org/.

##### Radial density analysis

To characterize morphology without connective branches, we performed a modified Sholl analysis^74^ by quantifying the distribution of Euclidean distances between each VT match and its soma. Point estimates were visualized using seaborn.kdeplot. Confidence intervals were generated by bootstrap resampling (1,000 iterations) and iteratively re-computing kernel density estimates (sklearn.neighbors.KernelDensity). To account for morphological differences^53,54^, apical but not basal matches within the soma layer were excluded. Comparative density across subfields (CA1, CA2, CA3) was calculated as the proportion of total radial density attributed to each group at a given distance.

##### Assessing superficial-deep heterogeneity

To quantify transcriptional differences between superficial and deep neurons, the soma layer was computationally bisected using cgrid. Differential expression was run with the limma implementation of the Wilcoxon Rank Sum test (logfc.threshold=0), and significant (Bonferroni-adjusted p-value <0.05) DE genes were bolded if also significant with a concordant sign logFC in published reference datasets^1,48^. Gene set enrichment analysis was performed using fgsea v1.24.0 on ordered (-log_10_(raw p-values)) protein-coding DE genes with scoreType “pos” for a one-tailed test against Gene Ontology Cellular Component terms (size 15–300), and hierarchically clustered (“ward.D2”) by Wang semantic similarity^63^ using the GOSemSim v2.24.0 R package^75^. Terms with leading edge of > 40 genes were visualized.

To account for the unbranched apical “neck” of deep pyramidal neurons^53,54^ when quantifying morphological differences (Fig. 6d-e), we computed the radial density distributions from each VT tree’s intersection point with the apical soma layer boundary, rather than from the soma itself. Analysis was restricted to VTs 40–400 μm from this boundary to exclude nuclear/cytoplasmic contamination and distal edge effects. Differences between superficial and deep distributions were quantified using the Wasserstein distance, with standard errors estimated via bootstrap resampling.

### Statistical Analysis

Statistical analyses were performed in Python (v3.8-3.11) and R (v4.2.2). Microscopy data are presented as mean ± standard error of the mean (s.e.m.) and experimental groups were compared with an unpaired, two-sided Welch’s t-test (degrees of freedom computed by Welch–Satterthwaite approximation); full details in figure legends (Fig. 1, Extended Data Fig. 1) and Methods. Projection plots report nominal jackknifed p-values with significance cutoffs of 0.05 in all analyses. Permutation t-tests for difference in mean were applied in the striatal spatial analysis using scipy.stats.ttest_ind, with p-values computed from two-sided tests. Correlation between striatum projection location and cortical dissection site was computed using the scipy.stats.pearsonr two-sided hypothesis test. Postsynaptic differences in quantile tests were computed using bootstrap resampling; p-values are from two-sided hypothesis tests. Differential expression and gene set enrichment analyses were performed and presented as indicated throughout the Methods.

## Supplementary Tables

**Supplementary Table 1.** Correlation of L5 ET cortical position and gene expression.

**Supplementary Table 2.** Gene set enrichment analysis results of L5 ET spatially correlated genes.

**Supplementary Table 3.** Differential expression results of medullary-projecting and non-medullary projecting L5 ET cells.

**Supplementary Table 4.** Gene set enrichment analysis of differential expression results in Supplementary Table 3.

**Supplementary Table 5.** Hippocampal subfield markers ascertained in Slide-seq.

**Supplementary Table 6.** Superficial-deep axis markers for each hippocampal subfield.

**Supplementary Table 7.** Gene set enrichment analysis results of superficial-deep marker genes.

**Supplementary Table 8.** Oligonucleotide primers used in this study.

**Supplementary Table 9.** Cell class markers used to filter out doublet snRNA-seq profiles.

**Supplementary Table 10.** Presynaptic projection results, p-value table.

